# Early Neurodevelopmental Defects in Huntington’s Disease are Driven by Choroid Plexus Overgrowth and Altered Paracrine Signaling

**DOI:** 10.1101/2024.09.23.614496

**Authors:** Karolina Świtońska-Kurkowska, Jakub Kubiś, Joanna Delimata-Raczek, Bart Krist, Magda Surdyka, Żaneta Kalinowska-Pośka, Piotr Piasecki, Luiza Handschuh, Jan Podkowiński, Magdalena Rakoczy, Anna Samelak-Czajka, Michael Hayden, Nicholas S Caron, Maciej Figiel

**Author notes:** Corresponding author: Maciej Figiel, Ph.D., Institute of Bioorganic Chemistry Polish Academy of Sciences Noskowskiego 12/14, 61-704, Poznan Poland. equally contributed (Karolina Świtońska-Kurkowska, Jakub Kubiś).

## Abstract

Huntington’s disease (HD), especially juvenile-onset HD (JOHD), involves early neurodevelopmental pathogenesis alongside the gradual breakdown of the corticostriatal neural axis. To better understand this mechanism, we created fused dorsal–ventral forebrain organoids from induced pluripotent stem cells (iPSCs) from JOHD to mimic early corticostriatal interactions in the disease. We observed characteristic growth phenotypes in HD organoids and found that these phenotypes were influenced by paracrine signals from opposite regions (dorsal-ventral and ventral-dorsal). These phenotypes were only partially rescued by conditioning with medium in HD compared to control organoids. We also investigated humanized HD embryonic mouse forebrains at E13.5 to validate the phenotypes. Using single-cell RNA sequencing (scRNAseq) and immunofluorescence of the internal structure of HD organoids, we observed early neurodevelopmental signs, including stalling during the phase of increased progenitor cell growth, delayed neuron maturation, and disrupted patterning between pallial and subpallial regions. A consistent feature across *in vitro* and *in vivo* models was an abnormal expansion of transthyretin (TTR)-positive cells resembling choroid plexus (ChP), along with ectopic expression of neuronal factors in ChP and a reduction in populations expressing intermediate progenitor and interneuron markers. In mosaic organoids that combined healthy and JOHD tissues, the healthy environment helped reduce ChP overgrowth and restore normal progenitor and neuronal development. This suggests that some developmental defects caused by mutant HTT are reversible and influenced by non-cell-autonomous factors. Overall, these findings point to early ChP-related abnormalities as a key aspect of JOHD neurodevelopmental disturbance and propose the ChP–CSF environment as an important area for future mechanistic studies and potential therapies.

## 1. INTRODUCTION

Huntington’s disease (HD) is a fatal, dominantly inherited neurodegenerative disease that affects mainly the central nervous system, causing the loss of neurons in the striatum and cerebral cortex (Cudkowicz and Kowall, 1990; Reiner et al., 1988). Symptoms may occur years or even decades prior to diagnosis, and the disease often worsens as the number of CAG repeats in the huntingtin gene increases (Quarrell et al., 2013). In rare cases, particularly with more than 60 repeats, the disease can occur before the age of 20, known as juvenile HD (JHD) (Fusilli et al., 2018), progresses more rapidly, and has different manifestations than adult-onset HD.

Pre-manifest HD patients can develop cognitive, psychiatric, and motor impairments long before diagnosis (Paulsen et al., 2014; Quaid et al., 2017). In particular, the phenotype that occurs years before overt HD symptoms is enlarged ventricular volume in the brain (Hobbs et al., 2010; Jahanshahi et al., 2022). The ventricular enlargement in HD patients is accompanied by enlargement of the choroid plexus (ChP) and parasagittal dural space (PDS), a major CSF outflow pathway. HD patients also demonstrated reduced ChP perfusion, pointing to aberrant CSF production and flow in HD (Hett et al., 2026). These phenotypes were proportional to the increase in CAG repeat expansion and with worsening motor impairment (Hett et al., 2026). Juvenile HD, also demonstrates an accumulation of various brain defects such as obvious ventricular enlargement. The symptom is also characteristic for neurodevelopmental diseases such as autism spectrum disorders (ASD) where it occurs together with abnormalities of the choroid plexus (ChP) and the formation of cysts of ependymal origin in early brain development (Movsas et al., 2013). The ChP is an important physiological barrier and the principal source of cerebrospinal fluid (CSF) responsible for clearance of toxic products from the central nervous system (CNS) (Lehtinen et al., 2013; Redzic et al., 2005). However, ChP also takes part in brain development and produces a powerful secretome, a mixture of signaling morphogens such as BMPs, Wnts, and SHH; growth factors; and regulators (IGF-2, NT-3, Otx2), and releases it directly into the cerebrospinal fluid (Ghersi-Egea et al., 2018; Rodrigues et al., 2026). These secreted factors coordinate neural progenitor proliferation, migration and differentiation. On the other hand, the development of the choroid plexus epithelium also depends on signaling such as BMP induction of epithelial fate and canonical Wnt\beta-catenin activity to prevent neuronal transformation of ChP. In particular huntingtin (HTT), is responsible for establishing apical-basal cell polarity, regulats ciliary function and Wnt signaling influencing epithelial transformation (Godin et al., 2010b, 2010a).

In HD, loss of neostriatal volume and medium spiny neurons (MSN) goes along with pyramidal neuron loss in the neocortex (Vonsattel et al., 2011). The combination indicates complex HD pathogenesis, including various cellular and communication impairments between the dorsal (cortex) and ventral (striatum) regions, particularly during forebrain development (Rangel-Barajas and Rebec, 2016). Recently, the brain organoid approach (Lancaster et al., 2013) was used to investigate HD (Conforti et al., 2018) and recently the method was enriched with fused cerebral organoids which recreate the dorsal-ventral forebrain axis (Bagley et al., 2017). Since such assembloids can reproduce dorso-ventral forebrain origin, they are an ideal model for studying the interactions between the most important and HD-involved brain regions. Brain organoids originating from juvenile HD cells would be particularly useful for defining neurodevelopmental HD brain pathogenesis at several levels, such as the molecular mechanisms, cellular organization of brain and blood-brain barrier (BBB) and choroid plexus impairments.

In our work, we have established a dorso-ventral HD juvenile fused organoids system to define the complex neurodevelopmental impairments at the levels of molecular biomarkers, cellular differentiation and interactions, and structural changes in the HD brain. We showed that assembling the dorso-ventral axis by fusing HD organoids differentiated towards the cerebral cortex (CTX) and the striatum (STR) can reveal distinct early brain neurodevelopmental features in HD. We found that dorsal and ventral HD organoids can control their growth in the paracrine non-cell-autonomous manner and that the internal structure of the HD fused organoids has net proliferative characteristics and a large prevalence of stem cell signatures compared to healthy organoids. We demonstrate early choroid plexus (ChP) cell population development in HD brain organoids and HD mouse model embryos compared to WT. In the dorsal part of embryo brain the choroid plexus is accompanied by lack of ventral population. All these phenotypes are normalized when we construct mixed HD-WT mosaic organoids. Moreover the HD organoids and HD mouse embryonic brains show defects in positioning of neural cell which disturbes cooperation of dorsal and ventral regions in brain development. We propose a conceptual model in which ChP abnormalities and dysregulation of MGE cell populations are essential neurodevelopmental features of HD pathogenesis. Future implications of our work point to ChP as a possible new direction in the search for therapeutic options, and to TTR as a very early developmental marker to monitor HD at the embryonic stage of the organism.

## 2. RESULTS

The experimental scheme of the entire manuscript is depicted in Fig. 1A,B,C,D,E,F. The culture system for the generation of fused dorso-ventral HD and control forebrain organoids, and experimental design is depicted in **Fig. 1A**. Control and HD dorso-ventral forebrain organoids were generated from human control 21Q (ND42245), 17Q/18Q (ND38555), 33Q (ND41113) and juvenile HD iPSC lines 71Q (ND42230), 77Q (CS77iHD-77nxx) and 109Q (ND42224) lines producing either single dorsal or ventral, fused control or HD and mosaic organoids. In brief, neurospheres were patterned into dorsal and ventral forebrain organoids, which were later fused and cultured to model early pseudotissues comprising the cerebral cortex (CTX) and the striatum (STR). Since we pursued modeling early brain development, we established day 60 as the main maturing endpoint for our control and HD organoids. However, for paracrine signaling, we evaluated the growth phenotype using organoids between 65 and 123 days (Fig 1B). Importantly, we also generated mosaic organoids, which served as a compensatory model to assess the contributions of dorsal and ventral regions to the development of HD phenotypes in fused organoids. The organoid names used at some points in the description of experimental procedures and results referred to the iPSC line names, the number of glutamines (Q), and the patterning into dorsal and ventral brain organoid regions. We produced the following organoids: 17Q Dorsal/17Q Ventral; 21Q Dorsal/21Q Ventral; 33Q Dorsal/33Q Ventral; 71Q Dorsal/71Q Ventral, 77Q Dorsal/77Q Ventral, 109Q Dorsal/109Q Ventral and mosaic organoids. Later in the manuscript, the names were abbreviated, such as: 17D/17V, 71D/71V, 71D/21V, and so on. The number indicates the number of Q or CAGs in the HTT protein or *HTT* gene.

**Figure 1.**
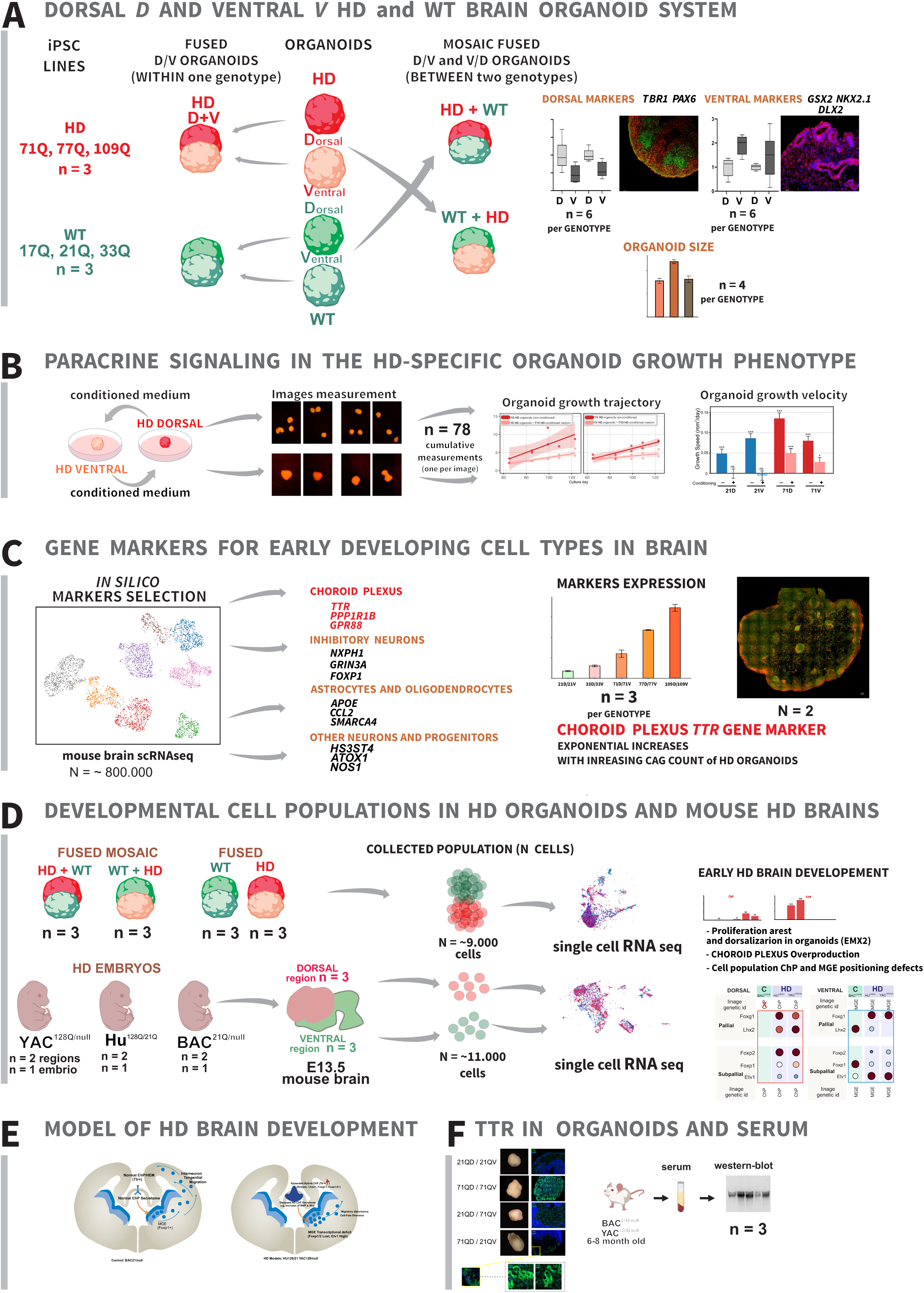
The scheme of the culture system for the generation of fused dorso- ventral HD and control forebrain organoids, and experimental design. The scheme of A) new model organoid system derived from n = 3 patients iPSC (genotypes) with indicated CAG counts. The system consists of separately generated dorsal and ventral organoids, by incubating neurospheres with dorsal or ventral small molecule primers, SHH inhibition by cyclopamine A for dorsal (mimicking cortex), inhibition of WNT by IWP2 and activation of SHH by SAG for ventral (mimicking striatum) brain patterning. The organoids were later fused into WT, HD, and WT/HD mosaic organoids (D/V and V/D configuration) aimed to observe developmental phenotypes and their rescue. The organoids were characterized by qPCR (n = 6 per marker) and immunofluorescence. B) The scheme of medium conditioning to understand how secreted factors can modulate development in a fused organoid model by using medium from single dorsal or ventral organoids both WT 21Q and HD 71Q. To determine non-cell-autonomous effects we used the trajectory and velocity of organoid growth starting on day 65 up to day 123. C) In the next step, the identification of gene markers of early developing cell types in the brain that included choroid polexus cells; inhibitory and excitatory neurons; and astrocytes and oligodendrocytes. The markers were first identified in silico for several dorsal and ventral brain regions using a population of N = 800,000 cells in total from public scRNAseq data, and a custom-made machine learning CSSG algorithm for scRNA analysis. The level of markers mRNA (qPCR, n = 3 organoids; immunofluorescence, n = 2-3 organoids) expression revealed the relative abundance of specific cell types. D) Detailed investigation of developmental neural cell populations affected by HD and rescue of identified population changes. The scRNAseq was performed in organoids (n = 3 per organoid type: WT, HD, and HD/WT mosaic; total of n = 12) and HD mouse E13 developing brains of Hu^128/21^ ^(null/null^ ^mouse^ ^BG)^ containing both human HD and normal allele, and containing only human HD or WT allele (YAC^128Q/null^, BAC^21Q/null^). n = 1 per genotype in total n = 3 and n = 1 per dorsal and n = 1 per ventral brain region in total n = 6 samples for scRNAseq per all regions and genotypes. The examination of specific populations and how positioning of cells in HD neural tissue during development occurred. E) We also constructed a conceptual model that may explain the observed neurodevelopmental phenotypes in HD which discussed in more detail in Figure 10. F) The analysis of the level of TTR marker of choroid plexus in organoids by immunocytochemistry. In addition, case study in HD animal sera derived from n = 3 Hu^128/21^ ^(null/null^ ^mouse^ ^BG)^ and n = 3 BAC^21Q/null^ mouse models were examined for TTR. To facilitate readability, the n-value (number of biological replicates) and N-value (cell population count) for each experiment were also provided directly on the entire scheme.

### 2.1. Juvenile HD and WT dorsal, ventral, and fused brain organoids demonstrate respective markers of early developing cortex and striatum

In order to confirm effective differentiation towards dorsal and ventral lineages, we compared 60-day-old brain organoids primed by dorsal or ventral patterning factors (n = 6 per genotype) used in the differentiation protocol in both WT and HD organoids **(scheme Fig. 1A)**. Therefore, we assessed the expression of specific cortical **(Fig 2A)** and striatal markers (**Fig. 2B**) and found correct dorsal or ventral specification in each type of separately primed organoid. The dorsal-primed organoids predominantly differentiated to dorsal (*TBR1* and *PAX6*) markers (**Fig. 2A**) while ventral priming produced organoids predominantly expressing ventral markers (*NKX2.1*, *DLX2* and *GSX2*) (**Fig. 2B**) as demonstrated via qPCR method. Subsequently, to visualize the enrichment of the cerebral cortex and striatum parts in 60 day fused organoids we detected dorsal **(Fig. 2C)** and ventral **(Fig. 2D)** early markers by immunostaining (N = 1 per genotype). We detected immunostaining for PAX6, which is a marker of the ventricular zone (VZ) and subventricular zone (SVZ) of the developing neocortex, respectively, and TBR1 as a dorsal forebrain marker **(Fig. 2C)**. For examining ventral organoid parts, we detected the immunofluorescence staining of DLX2 and NKX2.1, markers of medial ganglionic eminence (MGE), and GSX2, a marker of lateral ganglionic eminence (LGE) **(Fig. 2D)**. We also detected relatively late markers of differentiated neurons such as NEFH and SMI-32, indicating expression of forebrain markers, as well as other neuronal markers (**Fig. 2C,D**). Altogether, these results confirm the successful generation of organoids, with the presence of dorsal and ventral markers in separate dorsal or ventral organoids and the respective regions of fused organoids. Moreover, we demonstrate that the system provides an early striato-cortical microenvironment for examining emerging HD-specific developmental abnormalities.

**Figure 2.**
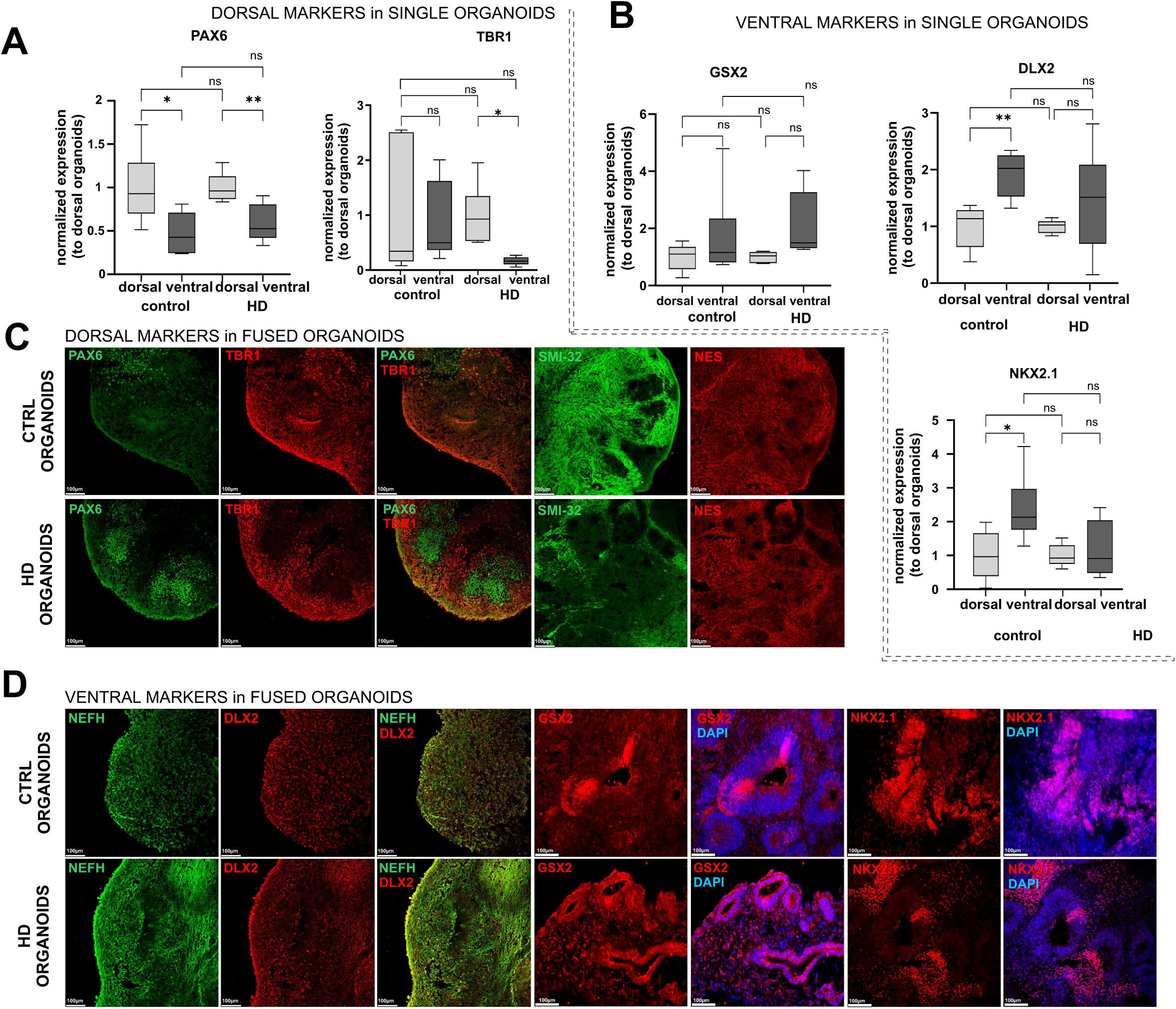
Juvenile HD and WT dorsal, ventral, and fused brain organoids show markers of early developing cortex and striatum. (A) qPCR for dorsal (*TBR1* and *PAX6*) and B) ventral (*DLX2, NKX2.1*, and *GSX2*) forebrain markers expression in 60-days-old of single dorsal and ventral HD and control organoids. C) Immunofluorescence of 60-day-old fused dorso-ventral HD and control organoids for cortical (TBR1 and PAX6) and D) striatal (DLX2, NKX2.1, and GSX2) markers, as well as for C) and D) NSCs (*NES*) and mature neurons *(SMI-32, NEFH*).

### 2.2. Fused juvenile HD brain organoids are large and contain prominent PAX6- neuronal rosettes, which are diminished in control and mosaic organoids

We examined the morphology and quantified the sizes (experimental scheme: Fig. 1A) of HD (n = 4 per genotype), mosaic (n = 4 per genotype), and control (n = 4 per genotype) organoids (**Fig. 3**). The 109Q HD organoids were significantly larger and were remarkably more complex in their shape than all control fused organoids. 60-day-old HD organoids were significantly larger in size (pval. < 3.35 e-05) than control organoids (**Fig. 3A and B**). On average, mosaic 21D/71V organoids were close in size to control organoids, while the opposite configuration of mosaic 71D/21V was close in size to HD organoids (**Fig. 3B**). All post-hoc test p-values are presented in **Fig. 3**.

**Figure 3.**
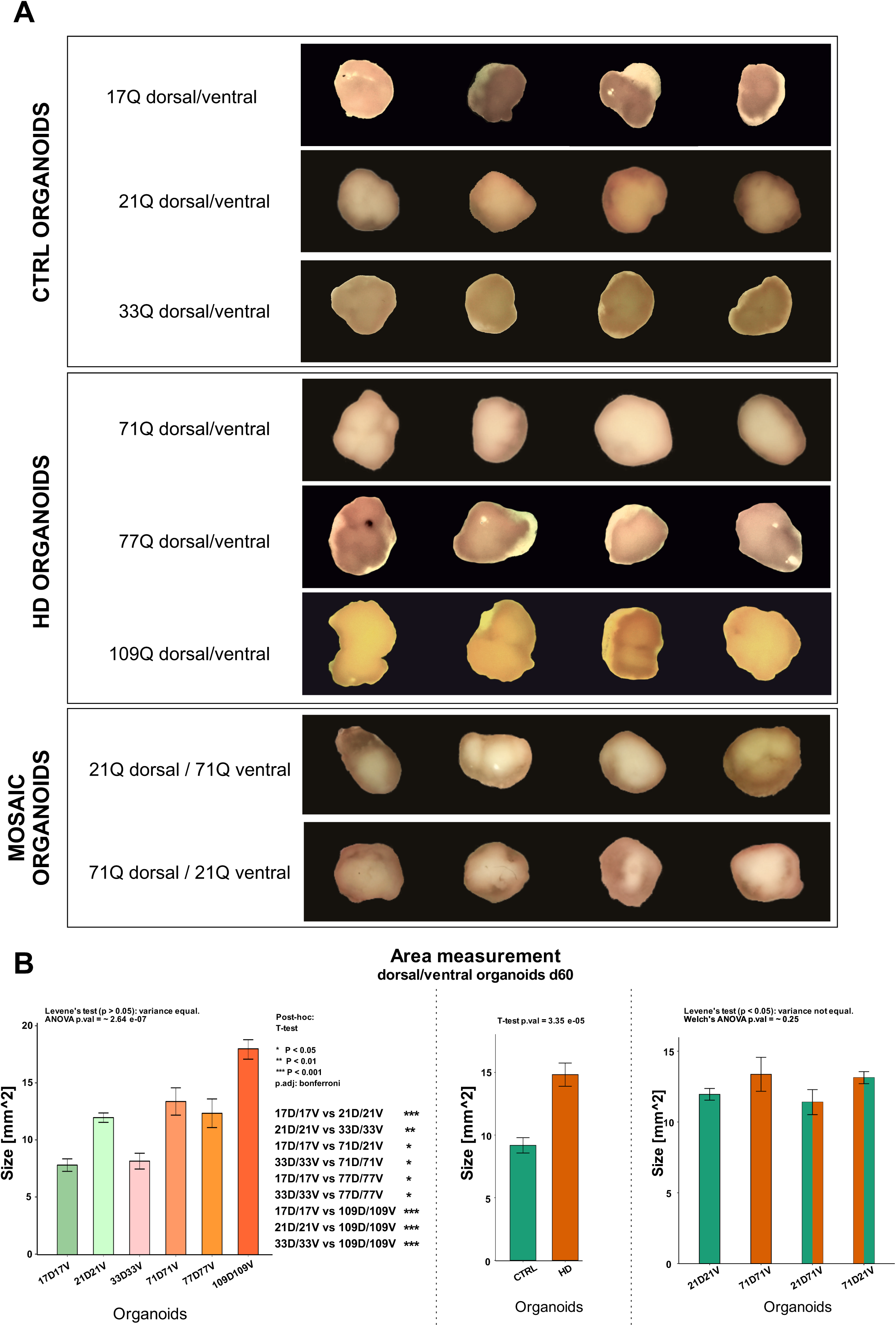
Macroscopic organization of 60-day-old HD fused organoids shows their less regular structure and increased size. A) Macroscopic 2D images of the fused dorsal/ventral control (17D/17V; 21D/21V; 33D/33V), HD organoids (71D/71V, 77D/77V,109D/109V), and mosaic HD/control organoids 71D/21V and 21D/71V; B) Area measurement graphs demonstrated data collected from 2D images of the organoids. The sizes of organoids derived from specific lines of the iPSC (n = 4 organoids per line) were compared to each other, and all stars of significance are depicted on the side of the main size graph. The sizes of dorsal/ventral fused HD organoids were significantly greater than those of controls. C) aggregated data panel (two bars) showing the mean size of all fused control vs all HD organoids. HD organoids were larger in size than controls. D) The sizes of the mosaic organoids vs control and HD organoids did not reach statistical significance (Welch’s ANOVA p=∼0.25); however, on average, 21D/71V was close in size to control organoids, and the opposite configuration of mosaic 71D/21V was close in size to HD organoids. The area [mm2] of each organoid was measured in ImageJ software. The following statistical tests were used: Levene’s test for examining data variance; if variance was equal, one-way ANOVA was used (p= ∼ 2.64e-07); post hoc: pairwise t tests with Bonferroni adjusted p values (p < 0.05, **p < 0.01, ***p < 0.001). If the variance was not equal, Welch’s ANOVA was used. The sample size: n = 4 per each organid type per iPSC line. Total n = 32.

In the developing brain, there is an expression of the *PAX6* marker, which indicates the presence of a population of early progenitors, and the expression of the *TBR1* marker, which indicates the presence of a population of early neurons. Our qPCR (**Fig. 2A**) and RNA-seq DEG results detected the *TBR1* [pval = 0.012387; log(FC) = ∼ 2.16] and *PAX6* [pval = 0.01252; log(FC) = ∼1.34] markers upregulation in 60-day dorsal brain HD organoids. To confirm the presence of the PAX6-positive and TBR-positive populations in our fused organoids, we performed immunostaining using the same type of organoids as used for scRNA-seq. Surprisingly, the immunostaining results on 60-day fused organoids (17D/17V n = 2, 21D/21V n = 2 (mutual CTRL n = 4), 71D/71V n = 2, 77D/77V n = 3 (mutual HD n = 5), and 21D/71V n = 2, 71D/21V n = 2 (mutual for mosaic n = 4) (**Fig.4A, B**) demonstrate the occurrence of a large number of PAX6-positive neural rosettes lying close to each other and the occurrence of the separate layers of TBR1 expression in HD fused organoids (71D/71V & 77D/77V). In contrast, the control fused organoids (17D/17V & 21D/21V) demonstrated only several scattered rosettes on organoid slices. Our results demonstrate that the development of the HD organoids is directed towards a more proliferative character of early neural development, with the occurrence of a vast number of progenitors in rosettes.

**Figure 4.**
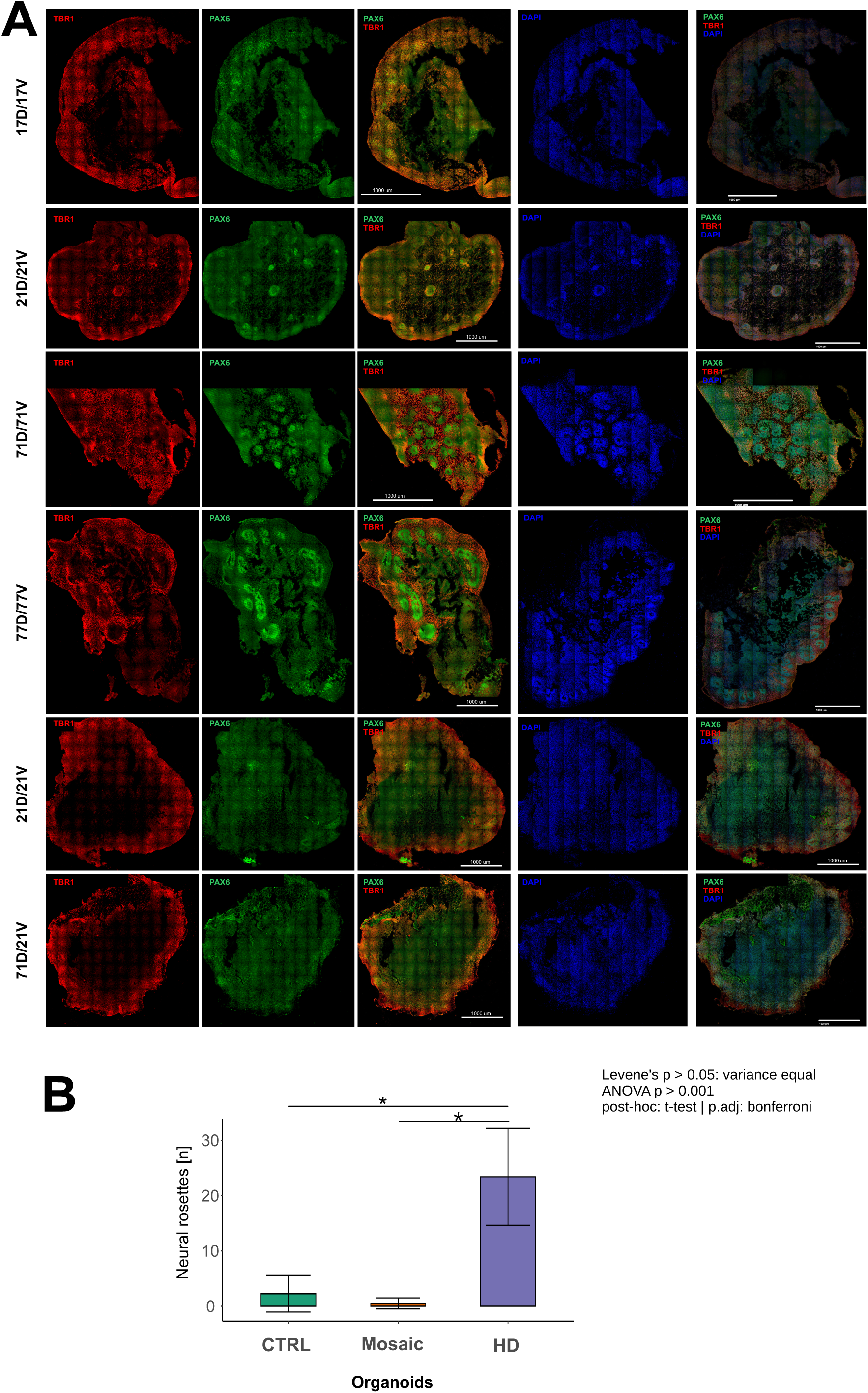
Prominent occurrence of neuronal rosettes containing TBR1 and PAX6- positive neural stem cell populations in 60-day-old fused dorso-ventral HD organoids. The internal structure of the organoids and the characteristic PAX6-positive, early progenitors, NSC) and TBR-positive, early-born neurons) dorsal populations in organoids were detected by immunostaining. Representative micrographs are labeled as HD (71D/71V, 77D/77V; n = 2 iPSC genotypes/HD patients), control (17D/17V, 21D/21V; n = 2 iPSC genotypes/normal patients), and mosaic organoid (21D/71V, 71D/21V) types. Each panel contains images of organoids co-stained with PAX6 (green) and TBR1 (red), as well as a merged image of both stainings for each organoid. Each panel also contains a DAPI stained image and a second merged image of DAPI/PAX6/TBR1 A) The micrographs revealed a large number of PAX6-positive neural rosettes lying close to each other and separate layers of TBR1 expression in HD fused organoids (71D/71V and 77D/77V), whereas in control fused organoids (17D/17V and 21D/21V), only several scattered rosettes and lower expression of PAX6 and TBR1 were present, indicating fewer neural stem cells and a less proliferative profile. In turn, HD organoids are directed toward the proliferative phase characteristic of early neural development, with a vast number of progenitors in rosettes. Importantly, in mosaic organoids, the HD phenotype becomes alleviated toward a more balanced proliferation. B) The panel demonstrates the quantification of roseetes showing the HD organoids show vast number of rosets wheres control organoids and mosaic have several rosetes.

Importantly, in the mosaic organoids, we observed the regression in the HD phenotype represented by reduced neural rosette occurrence **(Fig. 4A, B)**, which is consistent with the results from Over Representation Analysis (ORA) (**Supp. Fig. 1 A; Supp. 1**). Together, the enhanced proliferative architecture in HD organoids and regression in mosaic organoids suggest that early HD phenotypes can be mitigated through non-cell-autonomous interactions.

### 2.3. Paracrine signaling attenuates dorso-ventral growth without fully overriding the HD-specific organoid growth phenotype

The dorsal and ventral regions may interact through complex paracrine signaling. We wanted to understand how secreted factors can modulate development in a fused organoid model in both WT 21Q and HD 71Q subtypes. To determine whether such non-cell-autonomous effects occur in both HD and WT organoids, we used the trajectory of organoid growth starting on day 65 up to day 123 as a simple readout. We examined how dorsal or ventral organoids can influence each other’s growth by cultivating the organoids in medium containing 25% of conditioned medium within the genotypes (experimental scheme: Fig. 1B). For instance, we added 25% of ventral- or dorsal-conditioned media from HD organoids to their HD counterparts and compared conditioned versus unconditioned organoids within each genotype. To maintain stable nutrient and oxygen conditions and promote uniform molecular signaling within organoids, culture plates were kept on an orbital shaker at 60 RPM.

We revealed that altered growth was linked to organoid type (p=0.010), conditioning (p<0.001), and culture day (p<0.001) as well as interactions between conditioning and day (p<0.001), and group and day (p<0.001). Therefore, longitudinal growth was altered by both dorsal and ventral identity and conditioned medium. In general, all unconditioned organoids displayed steady longitudinal expansion, whereas the addition of conditioned medium suppressed growth **(Fig 5A, B)**. In addition, we compare the longitudinal organoid growth speed (in mm2 per day) across conditions. The analysis revealed continuous expansion in all unconditioned organoids (21D: 0.048 mm2/day, (p < 0.001); 21V: 0.086mm2/day, (p < 0.001); 71V: 0.080 mm2/day, (p < 0.001) and the expansion of dorsal HD 71 was over twice as fast: 0.135 mm2/day, (p < 0.001). Conditioning halted growth in WT organoids: 21D: -0.0006 mm2/day, (p = 0.958); 21V: -0.0055 mm2/day, (p = 0.631), whereas HD organoids exhibited reduced yet significant residual growth speeds 71D: 0.050 mm2/day, (p < 0.001); 71V: 0.027 mm2/day, (p = 0.021) **(Fig 5C)**.

**Figure 5.**
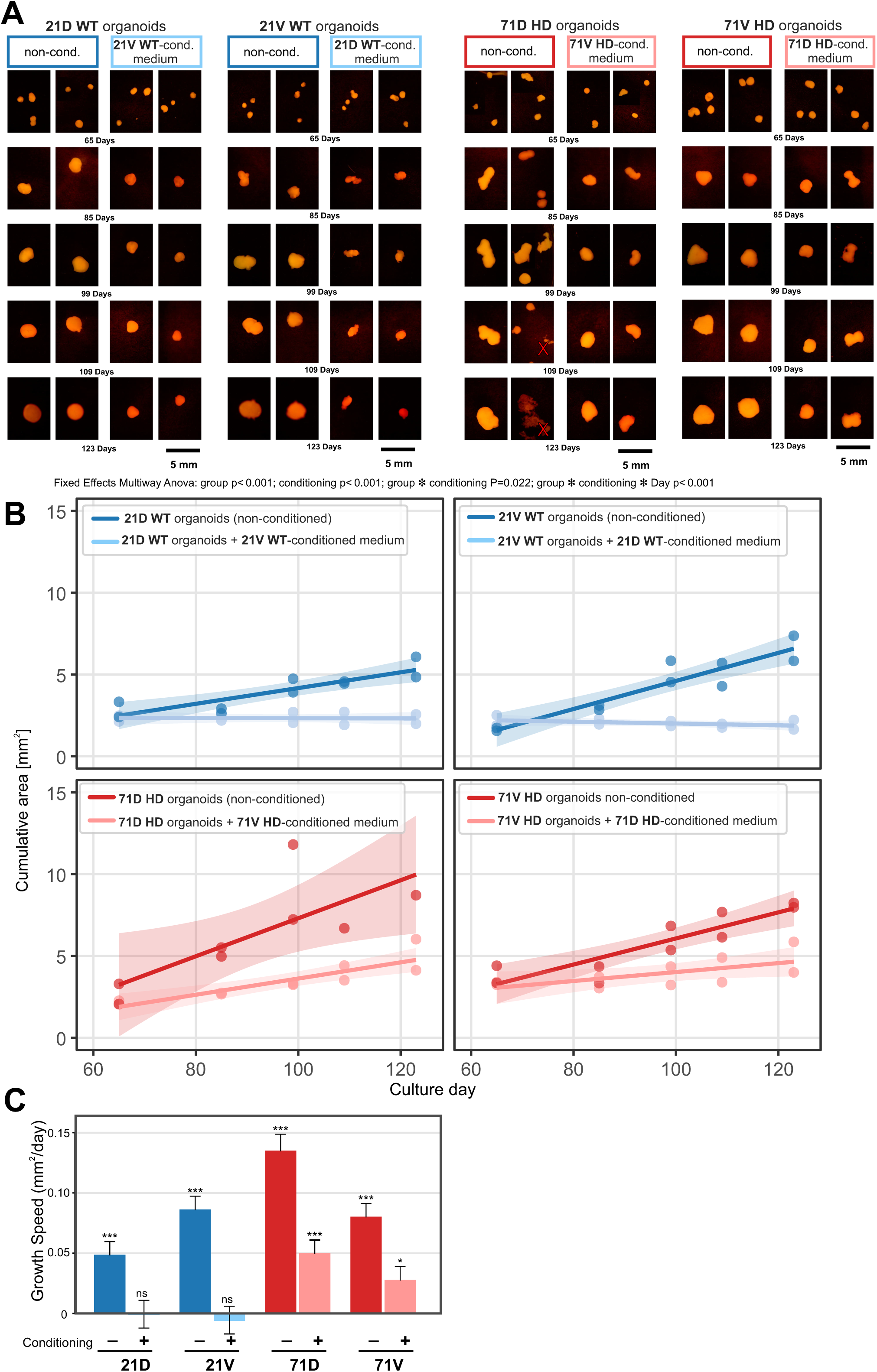
Investigation of paracrine interactions that control the development of organoid regions in a non-cell-autonomous manner. Dorsal or ventral and healthy or HD organoids were cultured with or without supplementation of a 25% medium conditioned in ventral or dorsal sites of the same genotype. In this paradigm, healthy 21 dorsal organoids were supplemented with 25% of medium conditioned by healthy 21 ventral organoids. Similarly, the following culture/supplementation configurations were set up: 21 ventral/21 dorsal and, in the case of Huntington’s Disease, 71 dorsal/71 ventral and 71 ventral/71 dorsal. As a simple readout, we used the trajectory of organoid growth by measuring the cumulative area (mm2) of organoids between day 65, when medium supplementation started, until day 123. A) The stereomicroscopic brightfield images of all conditioned and non-conditioned groups of healthy 21 and HD 71 ventral and dorsal organoids at day 65, 85, 99, 109, and 123. The organoids were measured from the collected images and subjected to statistical analysis. B) The graphs of cumulative surface area of healthy 21 dorsal and ventral and HD 71 dorsal or ventral organoids cultured alone (non-conditioned: healthy 21 dark blue and HD 71 red lines) or exposed to medium conditioned by the opposite region (conditioned: healthy 21 light blue and HD 71 pink lines). Statistical evaluation was performed via LMM and revealed that the differences were dependent on organoid type (p = 0.012), conditioning (p < 0.001), and day of treatment (p < 0.001). The differences observed were also affected by more complex interactions, such as the conditioning x day and group x day interactions (both p < 0.001). C) Moreover, we performed a simple slope analysis to compare the longitudinal organoid growth speed (in mm^2^ per day) across conditions. The analysis revealed continuous expansion in all unconditioned organoids (21D: 0.048 mm^2^/day, (***p < 0.001); 21V: 0.086mm^2^/day, (***p < 0.001); 71V: 0.080 mm^2^/day, (***p < 0.001) and the expansion of dorsal HD 71 was over twice as fast: 0.135 mm^2^/day, (***p < 0.001). Conditioning halted growth in WT organoids: 21D: -0.0006 mm^2^/day, (n.s. p = 0.958); 21V: -0.0055 mm^2^/day, (n.s. p = 0.631), whereas HD organoids exhibited reduced yet significant residual growth speeds 71D: 0.050 mm^2^/day, (***p < 0.001); 71V: 0.027 mm^2^/day, (*p = 0.021). This demonstrates that both dorsal and ventral organoid sides from both genotypes are subjected to regulatory developmental paracrine influence that controls tissue growth along the striatocortical axis and that HD is able to override this process despite paracrine influence. Data points represent individual well measurements, and lines indicate LMM- fitted linear trends. (Scale bar: 5 mm). Data n=78 are cumulative measurements derived from size observations, as the organoids were prone to aggregate over time. In most conditions and culture plates, 3 organoids were set up at day 65, which then aggregated into one, already by day 85. CI is 95% derived from the joint residual variance of the global LMM.

First, our experiment demonstrates pathological overgrowth of HD organoids vs WT organoids baseline where unconditioned 71 HD organoids grew larger and faster than unconditioned 21 WT controls between 65 and 123 day of organoid culture.

The paracrine modulation caused slightly distinct responces in dorsal vs ventral HD organoids. In dorsal HD organoids, supplementation of medium conditioned by 71V HD organoids attenuated the quick growth trajectory compared to untreated 71D controls. However, it was not able to completely suppres the intrinsic HD proliferative capacity, which may indicat that soluble paracrine signals alone are insufficient to override the HD phenotype. A growth reduction was also observed in ventral HD organoids (71V) upon exposure to 71D conditioned medium, indicating that ventral-dorsal secretome exchange in the HD background controls tissue expansion. Also, in the case of ventral HD organoids, the paracrine signals did not completely stop the growth. In contrast to HD organoids the paracrine signals from ventral or dorsal sides of healthy 21 organoids were able to completely prevent the growth of both 21D and 21V organoids compared to unconditioned healthy organoids.

These findings suggest that paracrine signaling between dorsal and ventral regions influences their development, affecting growth in both healthy and HD-related dorso- ventral patterns. However, the expansion of the HD organoid is twice as prominent compared to other types of organoids and can not be completely halted upon conditioning.

### 2.4. Prominent deregulations of cell population markers of choroid plexus, inhibitory neurons, and glial cells in juvenile HD organoids

We analyzed the publicly available scRNA-seq data (polpulation of N=∼ 800.000 cells; for experiment scheme Fig. 1C) *in silico* to select markers of the most common brain population of neuronal and non-neuronal cells for further analysis of their expression levels in dorso-ventral HD (n = 3 per genotype) and control organoids (n = 3 per genotype) (**Fig. 6**). Another step in the inclusion of markers for analysis was the manual selection, population markers likely affected in the developing HD brain. *TTR*, *H3ST4*, *PPP1R1B*, *GPR88*, *NOS1*, *NXPH1*, *APOE* and *GRIN3A* genes were selected. Additionally, using an *in silico* bioinformatic decision tree and *HTT* expression level approach (described in Materials and Methods), *ATOX1*, *FOXP1,* and *SMARCA4* were selected as marker genes for cell populations.

**Figure 6.**
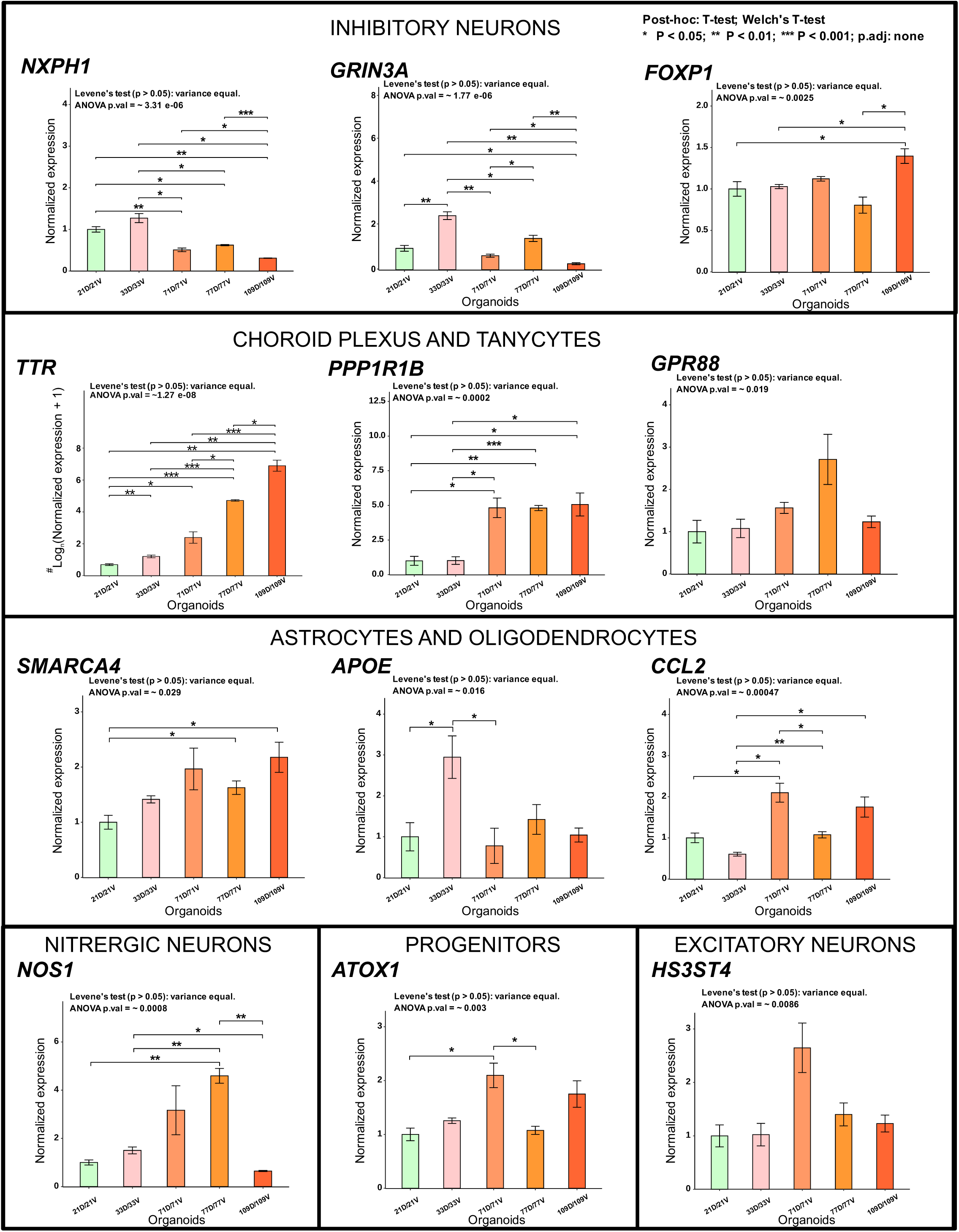
Evaluation of mRNA expression levels of neurodevelopmental marker genes in juvenile HD fused brain organoids. The panels are labelled with cell-type names and collect graphs showing the quantification of mRNA levels for marker genes in 60-day-old fused dorso-ventral HD and control organoids derived from several iPSC lines. The graphs are grouped by marker genes of inhibitory neurons (*NXPH1, FOXP1, GRIN3A*), choroid plexus (*TTR)* and tanycytes (*PPP1R1B, GPR88*), astrocytes and oligodendrocytes (*SMARCA4, APOE, CCL2*), nitrergic neurons (*NOS1*), progenitors (*ATOX1*), and excitatory neurons (*HS3ST4*). Particular attention was drawn to a highly upregulated and very statistically significant expression level of canonical choroid plexus marker gene *TTR* and tanycytes marker gene *PPP1R1B*. Moreover, *TTR* mRNA expression appears to increase gradually in organoids derived from iPSC lines with longer polyQ tracts. Moreover, there is a low level of NXPH1 mRNA, an interneuron marker, in all HD organoids, which may indicate low differentiation and migration of the cell type, suggesting halted cortex- and striatum-specific cell development. The mRNA expression levels of the genes in each type of fused organoid were compared with those of 21Q control organoids, which served as the basal expression level. The mRNA expression was normalized using *ACT*β as a reference gene. Levene’s test (p > 0.05) was used for variance examination, followed by one-way ANOVA (values indicated on the figure). Post-hoc Welch’s T-tests without p-value adjustment were used. The p values are indicated on the graph (*p < 0.05, **p 0.01, ***p < 0.001). The values: Mean ± SEM as error bars.

Selected genes were general markers associated with all basic types of brain cells, including inhibitory and excitatory neurons, astrocytes and oligodendrocytes, tanycytes, choroid plexus, nitrergic neurons, and neuronal progenitors.

Firstly, we compared the expression of several inhibitory neuron markers and showed the deregulated expression of 3 markers. *NXPH1* showed highly significant downregulation in all HD lines compared to 21D/21V and 33D/33V control organoids (ANOVA p.val = ∼ 3.31e-06). GRIN3A also showed downregulation while FOXP1 demonstrated inconsistent or non-significant outcomes **(Fig. 6)**.

Next, we analyzed the expression of selected choroid plexus (ChP) and tanycyte markers. *TTR* (ChP) and *PPP1R1B* (tanycytes) showed the most prominent upregulation in all HD lines compared to 21D/21V and 33D/33V control organoids. The changes in expression of the TTR mRNA were particularly differentiating across HD lines, with the increasing CAG lengths reaching 11.55 FC for 71D/71V, over 100 FC 77D/77V, and well over 1000 FC all vs 21D/21V control organoids. The 33D/33V organoids showed a 2.37 FC increase in TTR mRNA level vs. 21D/21V. The 33 CAG repeat line is still regarded as the healthy control; however, they contain a number of repeats that may indicate a certain risk of developing some HD phenotypes (**Fig 6**. ANOVA *p-value* ∼ 0.00036; posthoc: T-test * p<0.05, **p<0.01, ***p<0.001). Similarly, the tanycyte marker PPP1R1B showed a very consistent, high fold-change increase across all HD lines vs. control lines. DARPP-32, encoded by the *PPP1R1B* gene, plays a pivotal role in integrating signal transduction in dopaminoceptive neurons and has a regulatory role in the central nervous system (CNS) (Lin et al., 2021). Another tanycyte marker, *GPR88*, showed elevated expression in the HD line compared to 21D/21V and 33D/33V control organoids (ANOVA, *p-value* = 0.019). These results strongly point to the choroid plexus as the tissue most closely linked to HD pathogenesis.

In the group of astrocyte and oligodendrocyte markers, *SMARCA4* (ANOVA pval = 0.029) showed higher expression level in HD 77D/77V and 109D/109V, compared to 21D/21V control organoids, and *CCL2* (ANOVA pval = 0.00047) shows higher expression in HD vs 21D/21V or 33D/33V organoids. *ATOX1*, a marker of neuronal progenitors, showed increased expression in HD organoids (ANOVA pval = 0.003), while a post-hoc test showed a significant increase in expression in HD 71D/71V vs control 21D/21V organoids. HS3ST4, a marker of excitatory neurons (ANOVA = 0.0086), did not show a significant difference between organoid genotypes in the post-hoc test. CCL2 gene expression was increased in HD vs WT organoids (ANOVA = 0.0005), with post hoc tests showing significant differences between HD and 33D/33V and between 71D/71V and 21D/21V organoids. NOS1, a marker of nitrergic neurons, showed differential expression in HD vs WT organoids (ANOVA pval = 0.008), where post-hoc test showed a significant increase of expression between HD 77D/77V and 21D/21V or 33D/33V organoids. All post-hoc test p values are presented in **Fig. 6**.

Our results show selective changes in expression between fused control and HD organoids, indicating that mutant HTT alters lineages of cell populations, including inhibitory neurons, choroid plexus cells, and glial populations.

### 2.5. Early-developmental gene expression and ORA profiles in Juvenile HD organoids and the reversal in mosaic juvenile HD/Control

Data from scRNA-seq were first used to identify the total number of DEGs and determine the main processes involved in HD and development in our organoids and mosaic organoids. We pooled cell transcriptomes based on genotypes (pseudobulk method) and compared HD fused organoids (71D/71V, 77D/77V; n = 3 per genotype; total n = 6) vs. control (WT) fused organoids (17D/17V, 21D/21V; n = 3 per genotype; total n = 6), and mosaic HD/WT fused organoids (71D/21V, 21D/71V; n = 3 per genotype) vs. control fused organoids (21D/21V; n = 3).

Obtained DEGs are presented as volcano plots (**Fig. 7 A**). We obtained 7815 downregulated genes and 123 upregulated genes for HD (71D/71V, 77D/77V) vs. control (17D/17V, 21D/21V) fused organoids, 9038 downregulated genes and 177 upregulated genes for mosaic HD/WT 71D/21V vs. healthy (21D/21V) organoids, and 8141 downregulated genes and 49 upregulated genes for mosaic WT/HD 21D/71V vs. control 21D/21V organoids.

**Figure 7.**
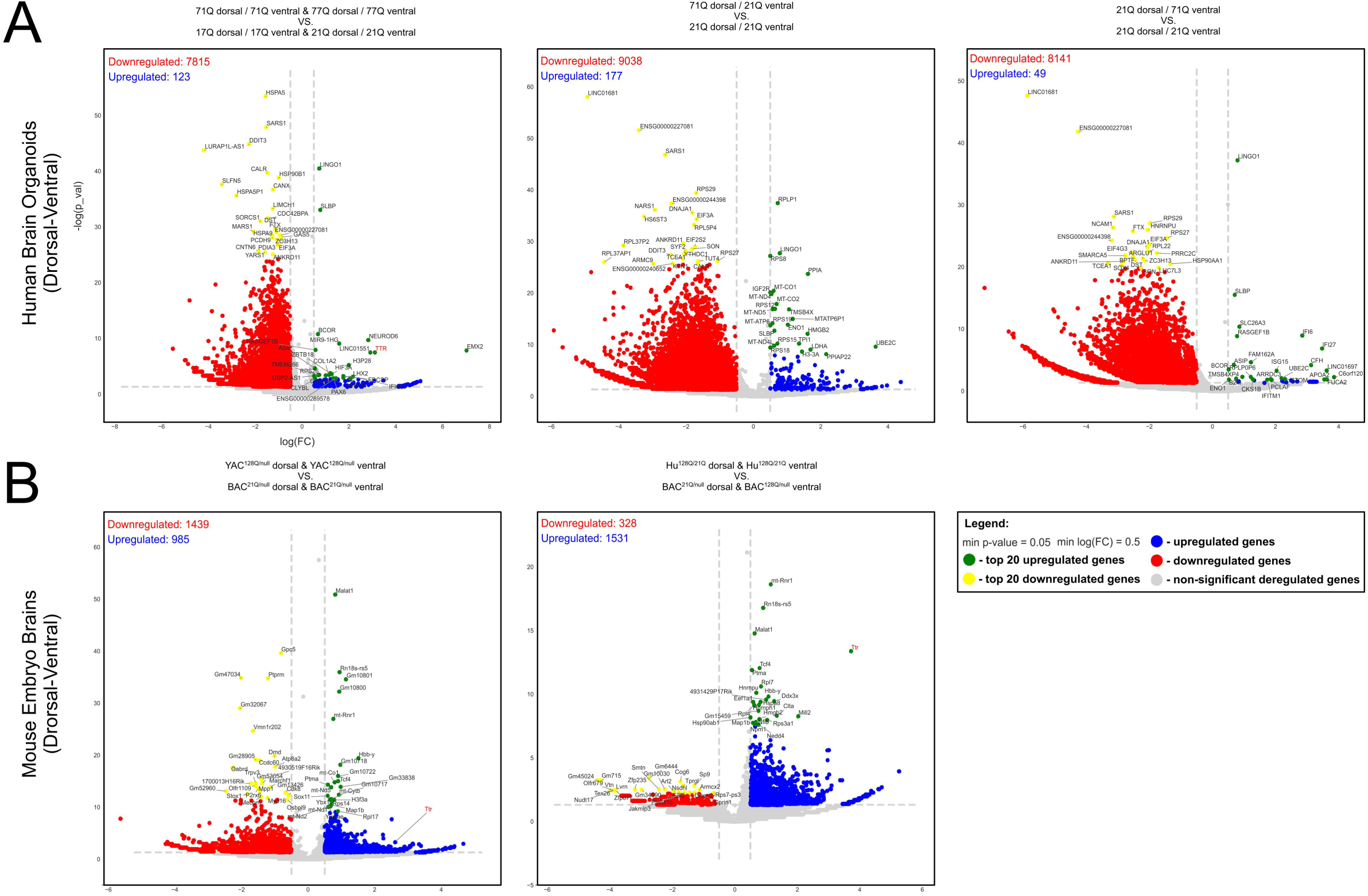
Identification of DEGs in transcriptomes of juvenile HD, control, and mosaic juvenile HD/control organoids, and HD embryonic brains. The volcano plots, demonstrate the DEGs identified in organoids and ventral or dorsal parts of HD embryonic brains, (YAC^128/null^ and Hu^128/21^) or control brains (Bac^21/null^). A) The analysis of DEGs for organoids consisted of the following comparisons of HD vs control organoids: 71D/71V & 77D/77V vs 21D/21V & 17D/17V, which have shown the most striking upregulation of the *TTR* expression in HD organoids, which is closely related to the occurrence of the ChP population. This deregulation was not significant either by comparing 71D/21V or 21D/71V mosaic organoid vs control 21D/21V organoids. B) The analysis of mouse embryonic brains has shown upregulation of the *TTR* in both models (YAC^128/null^ and Hu^128/21^). The analysis is extended by ORA (Gene Overrepresentation Analysis), available as supplementary figures 2 and 3. The statistical approach is described in the Statistics section.

Due to a certain variability of DEGs among organoids, we performed ORA (see Materials and Methods section 4.8). The terms that occurred most frequently across all organoids were manually assigned to arbitrary groups (broad functional headers on the supplementary figures) and presented in plots based on their p-values in the supplementary data (**Supp. Fig. 1, 2 & Supp. 1, 2, 3, 4, 5**).

In general, while terms enriched in both up- and downregulated datasets cluster under broad functional headers (such as ’Neuronal & glial cells development & maturation’), individual upregulated GO terms were specific to early progenitor maintenance and early differentiation steps, whereas downregulated GO terms were selectively enriched for late-stage synaptic assembly, dendritic spine formation, and mature neuronal/glial function.

The ORA demonstrated that HD organoids exhibit upregulation of genes associated with terms tightly connected to early development and brain formation (mostly Dentate gyrus, Forebrain, Telencephalon, and Eye), also connected with epigenetics and Extracellular Matrix (ECM) formation [*H3-3A, BIRC5, RPS26, PAX6, LHX2, NR2E1, EMX2, HMGA2, FEZF2, COL3A1, COL1A2, COL4A6, COL21A, VCAM1, TTR, LTBP3*]. In contrast, the terms enriched for downregulated genes in HD organoids are linked to processes characteristic of later stages of brain development, such as neuronal and glial cells maturation, migration, and signaling release related to processes of autophagy, mitotic spindle orientation and cilia motility. (**Supp. Fig. 1A**). The statistically significant deregulated genes between HD and CTRL organoids, along with the associated biological terms and pathways related to maturation, proliferation, and differentiation, indicate that cells derived from healthy organoids are more aligned with processes involving mature glial and neuronal cells. They also indicate that HD cells appear to be arrested in proliferative and differentiation stages. This is reflected in the faster development of certain brain structures in HD organoids, but with fewer processes linked to mature cells, suggesting an incomplete state of maturation. HD organoids delay or even arrest their development, preserving cell stemness, showing less relationship to terms indicating brain structure development compared to healthy organoids. It can also be closely linked to alterations in processes such as the mitotic spindle, cell polarity, cilia, cell motility, and autophagy, which play crucial roles in the early stages of healthy brain development. This is reflected by upregulated expression of specific gene markers such as *EMX2* [pval = ∼ 1,46E-08; log(FC) = ∼7.02] and *PAX6* [pval = ∼ 0,001; log(FC) = ∼1.34], which are early markers indicating a high degree of cell stemness in HD organoids. Moreover, the genes essential for proper cilia function in neurogenesis such as *IFT88* [pval = ∼ 4,58E-07; log(FC) = ∼-1.14], *ARL13B* [pval = ∼ 0.02; log(FC) = ∼-0.43], *KIF3A* [pval = ∼ 6,62E-06 ; log(FC) = ∼ -0.65], and mitotic spindle *MAPT* [pval = ∼ 4,99E-11; log(FC) = ∼ -1,2] are downregulated.

Furthermore, our ORA analysis revealed that several processes and terms observed for HD vs control organoids were not found or were reversed for mosaic organoids 71D/21V and 21D/71V vs. control organoids. For instance, terms characteristic of HD-upregulated genes related to initial brain formation were not found, and terms for HD-downregulated genes related to neuronal and glial maturation and migration were now identified for upregulated genes in mosaic organoids. (**Supp. Fig. 1 B & C; Supp. 1, 2, 3** for all genes, terms, and statistics).

The observation indicates that the interactions between ventral and dorsal parts in mosaic organoids can restore abnormal gene expression patterns associated with HD and normalize ORA signatures. This prompts an investigation into whether similar mechanisms occur *in vivo* during embryonic HD brain development.

### 2.6. Embryonic brains of HD mice contain gene expression and ORA profiles associated with HTT function

According the scheme from figure 1D, we compared mouse embryonic brains from HD Yac^128Q/null^ (one allele of human mutant HTT) vs control Bac^21Q/null^ (one allele of human normal HTT) and HD Hu^128Q/21Q^ (combination: 2 alleles, human normal and mutant HTT) vs control Bac^21Q/null^ (**Fig. 7 B**). We obtained 328 downregulated and 1531 upregulated DEGs for disease YAC^128Q/null^ vs. control BAC^21Q/null^, and 1439 downregulated and 985 upregulated DEGs for Hu^128Q/21Q^ vs. control BAC^21Q/null^ embryonic brains.

The analysis of YAC^128Q/null^ vs BAC^21Q/null^ demonstrated downregulated genes mainly related to neuronal and glial cell proliferation, development, and regulation of cell signal release, autophagy, and ECM formation. For upregulated genes, we noticed processes related to embryo development and brain structure formation, regulation of neurogenesis, cell migration, myelination, and cell connection formation and signaling related to processes of mitotic spindle orientation, cell polarity, cilia, and programmed cell death (**Supp. Fig. 2 A; Supp. 4**).

Similar results were observed in the Hu128Q/21Q vs Bac21Q/null analysis, except that downregulated genes also included terms and processes such as mitotic spindle orientation, cell polarity, and motility, and an increased number of genes were assigned to cell maturation and signal release terms. The upregulated genes demonstrated similar results to YAC^128Q/null^ vs BAC^21Q/null^ analysis (**Supp. Fig. 2 B; Supp. 5**).

In addition, results from both YAC^128Q/null^ and Hu^128Q/21Q^ mouse embryonic brains showed several upregulated genes involved in mitotic spindle regulation, potentially indicating altered development and neurogenesis in HD embryos. The upregulated genes included *Map1B* [YAC^128Q/null^: pval = ∼ 1,77E-08; log(FC) = ∼ 0.59, Hu^128Q/21Q^: pval = ∼ 3,47E-11; log(FC) = ∼0.69], *Clta* [YAC^128Q/null^: pval = ∼ 4,83E-09; log(FC) = ∼ 1.34, Hu^128Q/21Q^: pval = ∼ 0,00004; log(FC) = ∼ 1.02].

Together, all identified genes and ORA profiles indicate an excess of proliferative responses over differentiation responses connected with *HTT* function in the HD embryonic brain.

### 2.7. Increased TTR mRNA level is a common feature of HD organoids and HD embryonic brains compared to control, mosaic organoids, and control brains

Analyzing all our HD organoid and HD embryonic models, we found that *TTR* expression increased significantly in all HD models compared to their control counterparts. DEGs of 71D/71V & 77D/77V organoids showed greatly upregulated *TTR* expression log(FC) = ∼8.64 [pval = 3.41E-08] vs Control organoids (**Fig. 7 A**). Similarly the DEGs of YAC^128/null^ embryonic brain showed *TTR* expression log(FC) = ∼13.15 [pval = 6.03E-23], and Hu^128/21^ showed *TTR* expression log(FC) = ∼5.70 [pval = 2.1E-08] all vs control BAC^21/null^ embryonic brain (**Fig. 7 B**). Surprisingly, we found that our compensatory model, mosaic organoids in both HD ventral/control dorsal and control ventral/HD dorsal configuration, compared to the control 21D/21V organoids, did not reveal statistically significant changes in *TTR* expression (**Fig. 7 A**).

TTR protein often forms fibrillar deposits in the ECM in neurodegenerative diseases. Interestingly, the ECM formation genes consistently belonged to the downregulated gene pool in the HD embryo brain, and the same term was located in the upregulated gene pool in HD organoids. This could be attributed to the higher demand for extracellular matrix as a scaffold for cell proliferation in organoids, which are isolated model systems that are not as richly supplied with ECM as regular tissue, such as the brain. In the tissue, ECM demand is secured by various cell types and is more balanced.

### 2.8. HD organoids and embryonic brains contain a prominent increase in choroid plexus cells, which are not present or diminished in mosaic organoids

We performed droplet-based scRNA-seq (experimental scheme: Fig. 1D) across 3 experimental systems with multiple replicates per genotype: 1) HD and WT organoids, 2) mosaic HD/WT organoids, and 2) embryonic mouse brains containing either mutant human HTT, normal human HTT, or both and divided into ventral and dorsal parts.

In 60-day-old fused dorso-ventral JOHD (71D/71V and 77D/ 77V; [n = 3 per genotype]) and control (17D/17V and 21D/21V [n = 3 per genotype]) organoids where we profiled approximately N=8.000 single cells, and mosaic organoids (71D/21V and 21D/71V; [n = 3 per genotype]) with approximately N=3.000 single cells. In case of embryonic E13.5 mouse brains, we performed a n=6 scRNA-seq sequencing from: HD disease YAC^128Q/null^ [n = 1 per genotype including n = 1 dorsal brain and n = 1 ventral brain] and Hu^128Q/21Q^ [n = 1 per genotype including n = 1 dorsal brain and n = 1 ventral brain], and control BAC^21Q/null^ [n = 1 per genotype including n = 1 dorsal brain and n = 1 ventral brain] model where we profiled N=9.000 single cells.

We distinguished nine general cell populations across all analyzed organoids (**Fig. 8; Supp. Fig. 5**) and embryonic brains (**Fig. 9; Supp. Fig. 6**), characteristic of STR, CTX, MGE, LGE, caudal ganglionic eminence (CGE), hippocampus (HIP), MEL, and ChP. Clusters were annotated to reveal population identities as described previously (Kubiś and Figiel, 2023). This allowed us to “zoom in” on our results, divide the general populations into multiple subpopulations, and establish the cell lineage composition of the juvenile HD organoids and control organoids (**Supp. Fig. 3**) and of the mouse embryonic HD and normal dorsal and ventral parts of the brains (**Supp. Fig. 4**).

**Figure 8.**
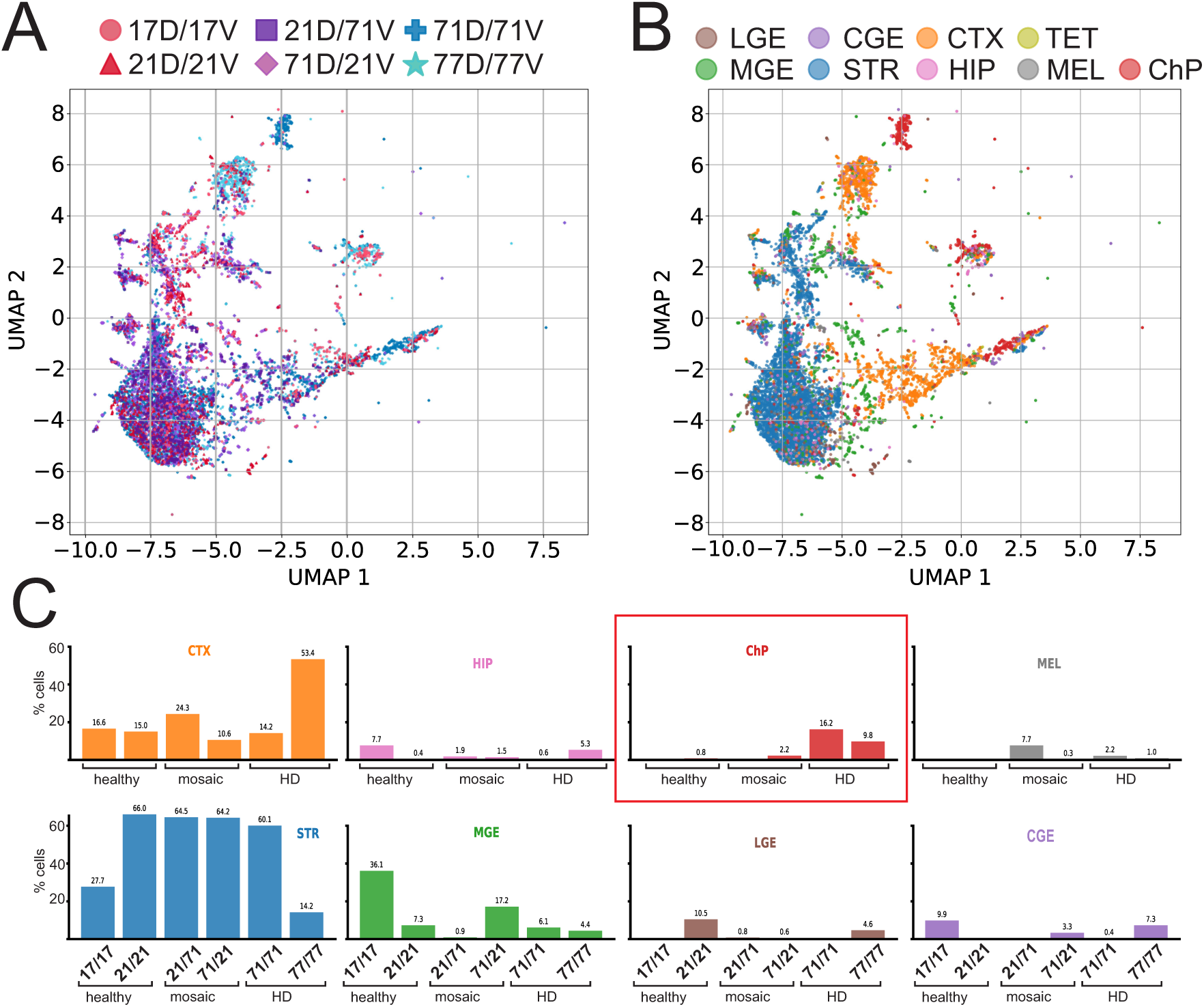
Composition of cell populations in HD (71D/71V; 77D/77V), control (17D/17V; 21D/21V), and mosaic (21D/71V; 71D/21V) organoids. The diagrams demonstrate scRNAseq data derived from organoids. A) To compare and visualize, cells from all six datasets HD (71D/71V; 77D/77V), control (21D/21V; 17D/17V), and mosaic (71D/21V; 21D/71V) were combined on a single harmonized UMAP graph, with each dataset/genotype, having n = 3; datasets, in total, n = 18 datasets). B) The main cell populations in HD and control organoids include dorsal regions: CTX (orange), HIP (pink), ChP (red), MEL (grey), and ventral regions: MGE (green), LGE (brown), CGE (violet), STR (blue), visualized on one harmonized UMAP. C) Bar charts illustrate the proportions of main brain region cell populations as percentages of all cells per organoid. Notably, the ChP population (red bars) is highly prominent in HD organoids, with some presence in mosaic organoids of the 71D/21V type. In control and 21D/71V organoids, ChP cells are rare or absent. The ChP population peaks in HD (71D/71V; 77D/77V) at 16% and 9.8%, respectively, and in mosaic organoids (71D/21V) at 2.24%. In contrast, control organoids (21D/21V) contain less than 1% ChP, and it is entirely absent in 17D/17V control organoids.

**Figure 9.**
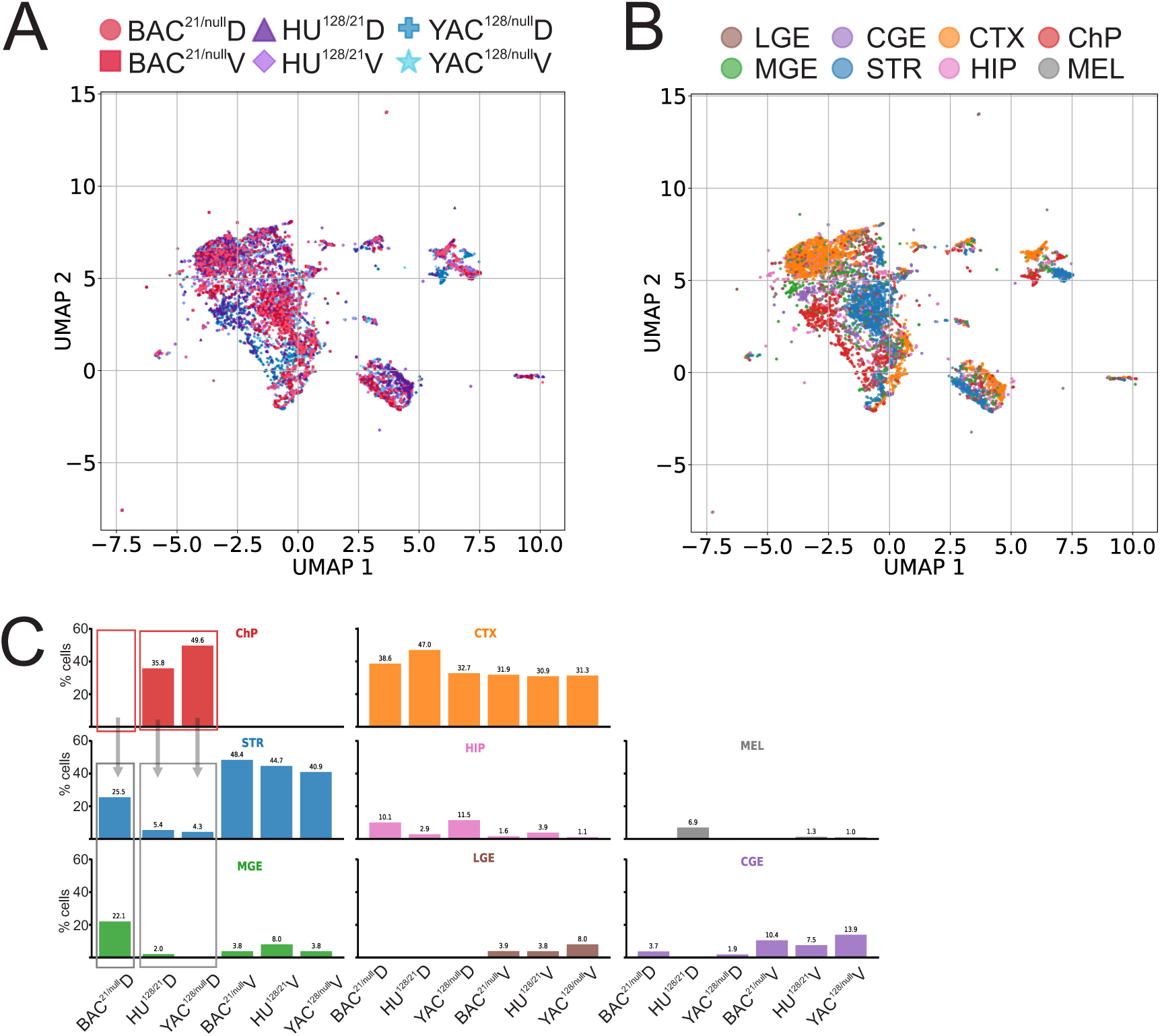
Composition of cell populations in dorsal and ventral brain parts of E13.5 embryos with mutant HTT (YAC128/null), normal and mutant (Hu128/21), and control (BAC21/null). scRNA-seq data were obtained from manually dissected ventral and dorsal regions of E13.5 embryonic brains in HD and WT mice. In addition to the ventral/dorsal separation, the tissue and resulting datasets differ in the composition of expressed human HTT, with no mouse HTT expression detected. (A) For comparison and visualization, all six scRNA-seq datasets from ventral or dorsal regions were integrated into a single harmonized UMAP plot. These datasets included dorsal and ventral samples from BAC^21/null^, YAC^128/null^, Hu^128/21^ genotypes (n=6 total: one embryo per genotype; two regions per embryo: dorsal and ventral). (B) Major cell populations shown on the UMAP included dorsal: CTX (orange), HIP (pink), ChP (red), MEL (grey); and ventral: MGE (green), LGE (brown), CGE (violet), STR (blue). (C) Bar charts illustrate the proportion of main brain region cell types, expressed as percentages of all cells from ventral or dorsal regions per genotype. Notably, the ChP population was prominent only in the E13.5 HD dorsal brain (YAC^128/null^ – 32.7%, Hu^128/21^ – 35.8%), and was absent in the control BAC^21/null^ dorsal brain. The cell mapping also revealed a pattern suggesting functional divergence: control BAC^21/null^ brains exhibited not only no ChP but also a significant population of ventral-derived cells MGE, migrating interneurons **(Supp.** Fig. 4). In contrast, HD embryonic brains (YAC^128/null^ and Hu^128/21^) showed near-complete absence of these populations and widespread ChP expansion.

71D/71V and 77D/77V HD fused organoids were highly populated with the TTR-positive choroid plexus cells, which accounted for as much as 10% and 16% of all cells. The ChP cells were also characterized by the expression of *TJP1* (pval = ∼ 2,54E-08; log(FC) = ∼7.59) and *GATA3* transcription factor (pval = ∼ 1,93E-05; log(FC) = ∼ 4.71), which is prominently active in the development and the *S100B* (pval = ∼ 0.00048; log(FC) = ∼ 3.13) (**Fig. 8**). Analysis of scRNAseq data showed that there was no ChP population in the 17D/17QV organoids and less than 1% in the 21D/21V control organoids. Another cellular population, present only in HD organoids, was melanocytes, which accounted for more than 2% and 4% in 71D/71V and 77D/77V fused organoids, respectively. The mosaic 71D/21V organoids, where the ventral side is healthy, we noticed that the contribution of ventral part cells (MGE [17.19%], CGE [3.33%], LGE [0.6%]) is higher than dorsal cells (CTX [10.63%], ChP [2.24%], HIP [1.49%]). The ratio of dorsal to ventral cells was opposite in mosaic 21D/71V organoids, where the dorsal side was healthy, and the ventral side was affected. In this case, the share of dorsal part cells (CTX [24.32%], MEL [7.67%], HIP [1.86%]) is higher than that of early ventral cells (MGE [0.88%], LGE [0.77%]). In both mosaic organoids, the proportion of STR ventral cells was the same, ∼64%. This demonstrates the imbalance in the development of the ventral and dorsal HD cell types in mosaic organoids.

Together, in HD organoids, more cells are directed toward cortical/dorsal patterns, including overproduction of ChP, compared with lower production of ventral-patterned cells. In mosaic organoids, the overproduction of *TTR*-positive ChP was rescued or attenuated, depending on whether the dorsal or ventral part was healthy.

Similar to HD organoids, ChP populations accounted for a large fraction of the dorsal embryonic HD brain in the YAC^128Q/null^ (49.64%) and Hu^128Q/21Q^ (35.78%), among all cells analyzed (**Fig. 9**). Embryo brains BAC^21Q/null^ control lacked ChP populations. In Hu^128Q/21Q^. Furthermore, thanks to the precise dissection of the embryo brain into ventral and dorsal parts (e.g. absence of ventral derived CGE, LGE on the dorsal site; also see section 2.10), the mapping of cell populations revealed a pattern pointing at functional divergence between control and HD embryo brains. Control BAC^21Q/null^ embryonic brains, in addition to the lack of ChP, exhibited a prominent, well-structured population of ventral-derived *Sox6*+ cells (MGE). Such populations of MGE *Sox6*+: MGE Sox6 Plxna4 – Cdh13; MGE Sox6 Plxna4 – Mmp16; MGE Sox6 Plxna4 – Csmd1; MGE Sox6 Plxna4 – Nnat, MGE Sox6 Plxna4 – Dcc; MGE Sox6 Plxna4 – Gm816; MGE Sox6 Plxna4 – Plxna4; MGE Sox6 Plxna4 – Dbx2; MGE Sox6 Gm10719 – Gm33838; MGE Sox6 Garem1 – Prkca; MGE Sox6 Garem1 – Gm48228; MGE Etv1 Plxna4 – Fmnl2 **(Supp. Fig. 4)** in the dorsal brain are predominantly attributed to migrating interneurons streaming from the MGE toward the dorsal telencephalon (Azim et al., 2009; Batista-Brito et al., 2009). On the contrary HD embryonic brains (YAC^128/null^ and Hu^128/21^), dominated by massive ChP expansion, showed a depletion or complete absence of migrating interneuron populations.

These scRNA-seq results from *in vivo* HD embryos are highly convergent, showing that the dorsal brain region overproduces choroid plexus cells, which restricts ventral populations in the dorsal telencephalon, strongly supporting a mechanism of HD pathogenesis based on microenvironmental disturbance and interactions between the cortex, ganglionic eminences, and striatum.

### 2.9. Disorganization of dorsal-ventral cell positioning in HD and its partial rescued in mosaic organoids

To evaluate the accuracy of dorsal-ventral (pallial-subpallial) positional identity in our models, we used scRNA-seq to analyze the composition of cell lineages in HD organoids (experimental scheme **Fig. 1D,E**). Specifically, we have selected *EMX2*, *LHX2*, and *FOXG1* as telencephalic markers characteristic of early neuronal stem cells and progenitors, which are primarily localized in the developing cortex and are also expressed in ganglionic eminences to some extent. As markers of subpallial cell positioning, we have selected *FOXP1* and *FOXP2*.

In healthy control organoids, cells within the cortical (CTX) and hippocampal (HIP) clusters showed strong expression of the mature deep-layer neuronal marker *TBR1* and the cortical selector *LHX2*. Meanwhile, the early progenitor factor *EMX2* was almost completely absent. This pattern indicates that the control cells successfully exited the early proliferative stem cell stage and moved into later stages of cortical neurogenesis. This molecular profile matches our overall observations, where the healthy control organoids appeared solid and exhibited a balanced growth rate.

In contrast, pure HD mutant organoids continued to exhibit high, sustained levels of *EMX2* and *LHX2* in their dorsal regions. This indicates a developmental delay, with cells staying in an early progenitor-like state and a significant halt in cortical maturation. Notably, the HD lines exhibited severe disruption of their positional identity: markers typically found in the upper brain, such as *EMX2* and *LHX2*, were abnormally expressed in cell populations showing ventral markers and assigned to ventral areas, particularly in the STR and MGE. Additionally, the cell populations assigned to the MGE still expressed the dorsal marker *FOXG1*. These molecular changes help explain the visible abnormalities, including overgrowth, a disorganized appearance of the HD organoids, and the overproliferation of rosette structures. **(Fig. 10A)**

**Figure 10.**
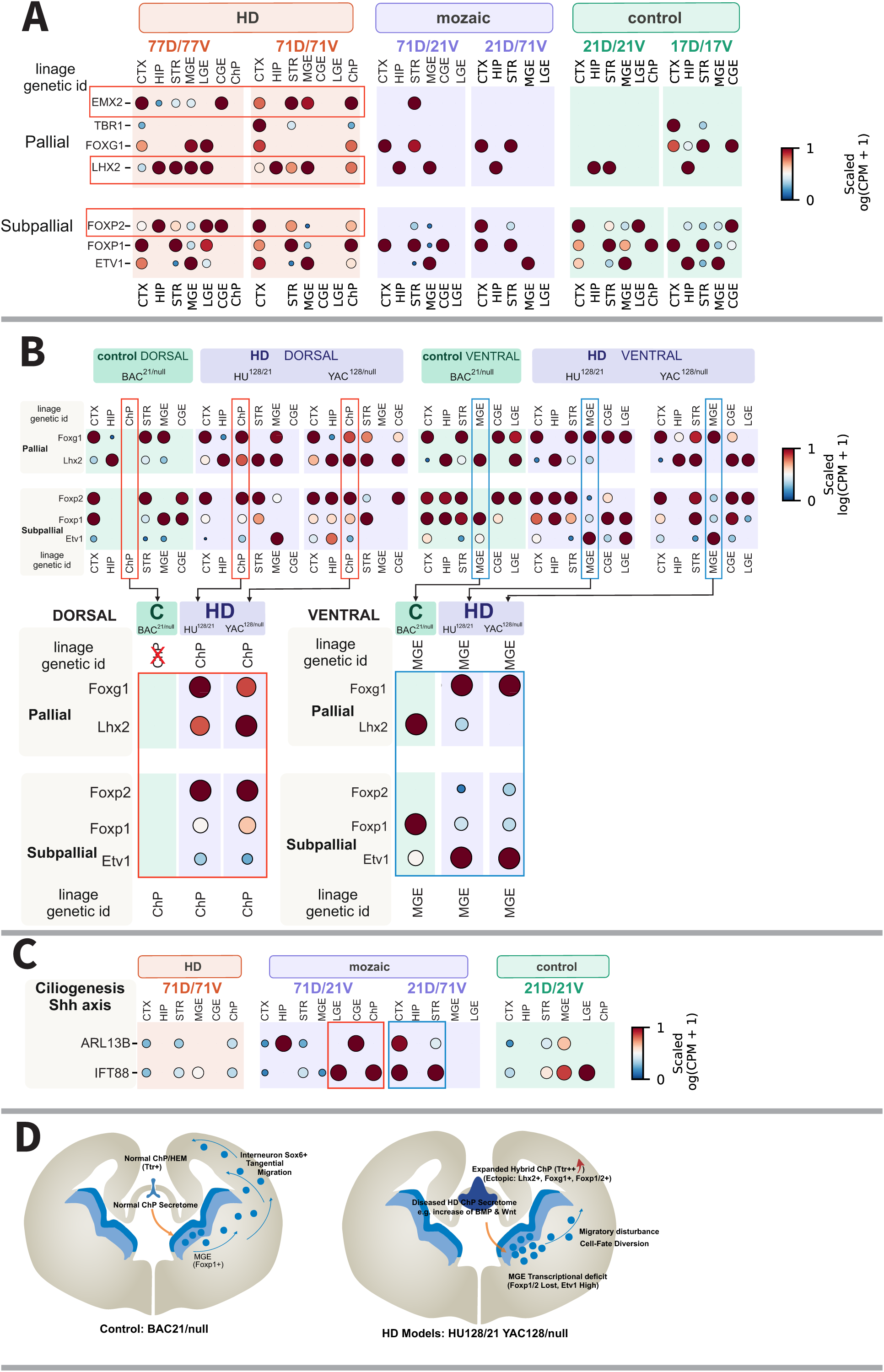
Huntington’s disease mutation disturbs pallial-subpallial lineage fidelity, resulting in ectopic ChP, MGE transcriptional dysregulation and non-cell autonomous regional dorsalization. (A) Loss of spatial identity and regional compartmentalization in HD fusion organoids. Dot plots of scRNA-seq data illustrate the expression profiles of pallial (FOXG1, EMX2, TBR1, LHX2) and subpallial (FOXP1, ETV1, FOXP2) lineage markers across distinct cell clusters: cortex (CTX), hippocampus (HIP), striatum (STR), medial ganglionic eminence (MGE), lateral ganglionic eminence (LGE), caudal ganglionic eminence (CGE), and choroid plexus (ChP). Healthy control organoid lines (21D/21V, 17D/17V) demonstrate proper restriction of positional dorsal markers in ventral structures such as MGE and CGE. Ventral markers are sparsely expressed in dorsal populations such as CTX and HIP, showing that the organoid model is not as faithful as the embryonic brain. HD mutant organoids (77D/77V, 71D/71V) show profound spatial confusion, marked by the ectopic expression of early progenitor dorsal markers (EMX2, LHX2) within the ventral STR and MGE populations, alongside anomalous leakage of FOXP2 into the CTX and HIP domains. Mosaic organoids integrating healthy and mutant cells (71D/21V, 21D/71V) reveal a selective rescue of this positional phenotype, actively halting the dorsalization cascade. B) Loss of lineage fidelity and specific MGE vulnerability in the embryonic HD mouse forebrain. (B) *In vivo* scRNA-seq analysis of dorsal and ventral telencephalic regions from E13.5 mouse embryos, comparing control (BAC21/null) to HD models (HU128/21, YAC128/null). The data visualize the spatial distribution of Foxg1, Foxp1, Etv1, Foxp2, and Lhx2 across CTX, HIP, STR, MGE, CGE, LGE, and ChP clusters. The highlighted raws with expression profiles reveal severe subpallial compartmentalization defects. While LGE and CGE clusters in HD embryos largely preserve Foxp1 and Foxp2 expression, the MGE cluster undergoes a targeted transcriptional collapse in HD embryos. This is defined by the loss of Foxp1 and Foxp2, accompanied by the pathological, ectopic activation of the differentiating neuronal marker Etv1 within the MGE. Importantly, the dorsal HD choroid plexus exhibits ectopic expression of Foxg1 and Lhx2. C) Alterations in ciliogenesis and Shh signaling pathways. Dot plots displaying the altered expression patterns of ciliogenesis-related genes, ARL13B and IFT88, across telencephalic clusters (CTX, HIP, STR, MGE, LGE, CGE, ChP) in pure HD (71D/71V) and mosaic (71D/21V, 21D/71V, 21D/21V) organoids. D) Conceptual mechanistic model of non-cell autonomous HD pathogenesis. A schematic illustrates the proposed signaling cascade disrupting forebrain development in HD. An expanded, hybrid choroid plexus (Ttr++) ectopically expressing Lhx2, Foxg1, and Foxp1/2 drives the increased secretion of morphogens, such as BMP and Wnt, compared to a normal ChP/HEM (Ttr+). This pathological secretome acts non-cell autonomously on the ventral subpallium, leading to MGE transcriptional collapse (Foxp1/2 loss, high Etv1), cell- fate diversion, and the subsequent arrest of tangential interneuron migration. Lack or insufficient interneuron support disrupts corticostriatal interactions.

This evidence of spatial disorganization is further visible on the panel showing the subpallial identity. In control organoids, *FOXP2* is visible in populations assigned to ventral subpallial regions such as the STR, MGE, LGE, and CGE, while in HD organoids, *FOXP2* is abnormally expressed in the CTX and HIP. Additionally, important markers such as *FOXP1* and *ETV1*, which are linked to the subpallium, appear irregularly distributed across both dorsal and ventral regions in the HD models. This suggests that the mutant HTT may cause ventral subpallial cells to acquire dorsal features, which disrupts proper tissue compartmentalization during early development.

Notably, mosaic co-cultures (71D/21V and 21D/71V) showed that introducing healthy cells (21 CAG) could correct this positional defect. In the MGE clusters within these mosaic organoids, the important marker *LHX2* stayed active, even though its downstream targets *EMX2* and *FOXG1* were turned off. Meanwhile, *FOXP1* and *ETV1* gradually regained their normal, compartmentalized expression patterns. This suggests that adding healthy wild-type cells to mutant ones can reduce dorsalization and promote more balanced tissue development, partially restoring regional boundaries **(Fig 10A)**. This non-cell-autonomous attenuation highlights the significant role of the microenvironment in counteracting positional defects in HD cells, a phenomenon further explored through scRNAseq in the HD embryonic forebrain.

### 2.10. Loss of Lineage Fidelity and Ganglionic Eminence Vulnerability in HD Embryonic Forebrain

Building upon the evidence of spatial disorganization and microenvironmental influence in our *in vitro* model, we next investigated whether similar disorganization in tissue compartmentalization occurs *in vivo* **(Fig. 1E)**. Using the advantage of separated (dissected) dorsal and ventral telencephalic regions, defined by populations identified from scRNAseq of E13.5 HD and control mouse embryos’ brains **(Fig. 9)**, we were able to evaluate the pallial and subpallial developmental lineages of the HD brain by studying the telencephalic *FoxG1* and *Lhx2* markers and the more ventral *Foxp1*, *Foxp2*, and the marker of differentiating ganglionic neurons, *Evt1*.

First, we evaluated the transcriptomes of cell populations from the ventral compartment to assess the robustness of segregation of subpallial structures and to rule out technical artifacts arising from dissection of the ventral and dorsal parts of embryonic brains **(Fig. 10B)**. The cell populations from control BAC21/null embryos recapitulated canonical gradients of *Foxp1* and *Foxp2* across the ganglionic eminences (Ferland et al., 2003; Takahashi et al., 2003). The LGE showed strong *Foxp2* expression levels but lacked *Foxp1*, while the MGE displayed the opposite pattern, with high *Foxp1* levels. The CGE populations had an intermediate profile, predominantly expressing *Foxp2* with moderate *Foxp1*. In the STR cluster, both *Foxp1* and *Foxp2* were consistently expressed across HD and controls, though *Foxp1* was slightly reduced in the heterozygous HU^128/21^ embryo. This consistent spatial pattern of core striatal and subpallial populations after scRNAseq suggests that our tissue dissection was very precise and that our dataset offers high resolution.

Interestingly, while the LGE and CGE population clusters in YAC^128/null^ and HU^128/21^ maintain stable and mostly unchanged levels of *Foxp1* and *Foxp2*, the MGE cluster shows a strong decrease in the expression of these markers and also a reduction in dorsal *Lhx2* in both HD models. In addition to the selective loss of the MGE gene expression program, we observed abnormal activation of *Etv1* within the MGE populations in HD. Normally, *Etv1* is a marker of postmitotic, maturing neural cells migrating from the lateral subpallium, and not a marker of early, proliferating progenitors. This abnormal profile suggests a disruption in the compartmentalization of the subpallium **(Fig 10B)**.

This disruption in gene expression within MGE interneuron progenitors does not occur in isolation but co-occurs with abnormally expanded ChP that likely generates the changed signaling secretome, suggesting a non-cell-autonomously driven effect. At the single-cell level, expanded choroid plexus cells in HD showed abnormal hybrid expression patterns that combined markers from both dorsal and ventral regions, indicating a failure in normal epithelial cell development. Within the subpallial MGE area, this molecular disorganization manifested to a significant decrease in the vital migratory transcription factor *Foxp1* and an early, abnormal activation of the postmitotic maturation marker *Etv1*. This switch in gene expression causes cells to exit the cell cycle prematurely and stop migrating, trapping interneurons within the MGE.

Together, these findings establish a conceptual model **(Fig. 10D)** where massive ChP overgrowth creates an altered paracrine environment that triggers the switch in gene expression in interneuron progenitors. In healthy conditions, *Foxp1*-positive interneuronal progenitors normally proliferate and migrate from the MGE. In HD, the overgrown choroid plexus influences premature differentiation into *Etv1*-positive neurons, which decreases interneuron generation and tangential migration and disrupts the structural assembly of the corticostriatal axis.

### 2.11. Choroid plexus structures in HD organoids and the presence of TTR Protein in the blood serum of the HD Mouse Model

We performed immunostaining (experimental scheme: Fig. 1F) for TTR in 60-days old fused dorso-ventral control 21D/21V (N = 1), HD 71D/71V (N = 1), and mosaic 21Q/71Q, 71D/21V (N = 1 per genotype) organoids (**Fig. 11A**). The results showed an abundant signal for TTR in 71D/71V HD, partial signal in 71D/21V mosaic organoid and no signal in 21QD/21Vorganoids, which corresponds to the scRNA-seq results. Transthyretin (TTR), also known as prealbumin, is a 55 kDa homotetrameric protein that transports thyroxine and retinol-binding protein (RBP) in the serum and cerebrospinal fluid. Since we found such a significant contribution of choroid plexus populations in HD organoids and HD embryonic brains and differential ocurance in mosaic fused organoids, we investigated the levels of TTR protein as a possible HD marker candidate in blood serum of adult 6-8 month old HD mice (**Fig. 11B**). Using Western blot assay, we investigated n = 5 YAC^128/null^ HD vs n = 5 BAC^21Q/null^ mouse sera and assessed both the level of TTR monomer (16 kDa) and tetramer (55 kDa). We showed that the marker differed between individual YAC^128/null^ mice and BAC^21Q/null^ controls (**Fig. 11**), mainly for the Ttr tetramer. We, however, did not observe statistical significance in the mean comparisons.

**Figure. 11.**
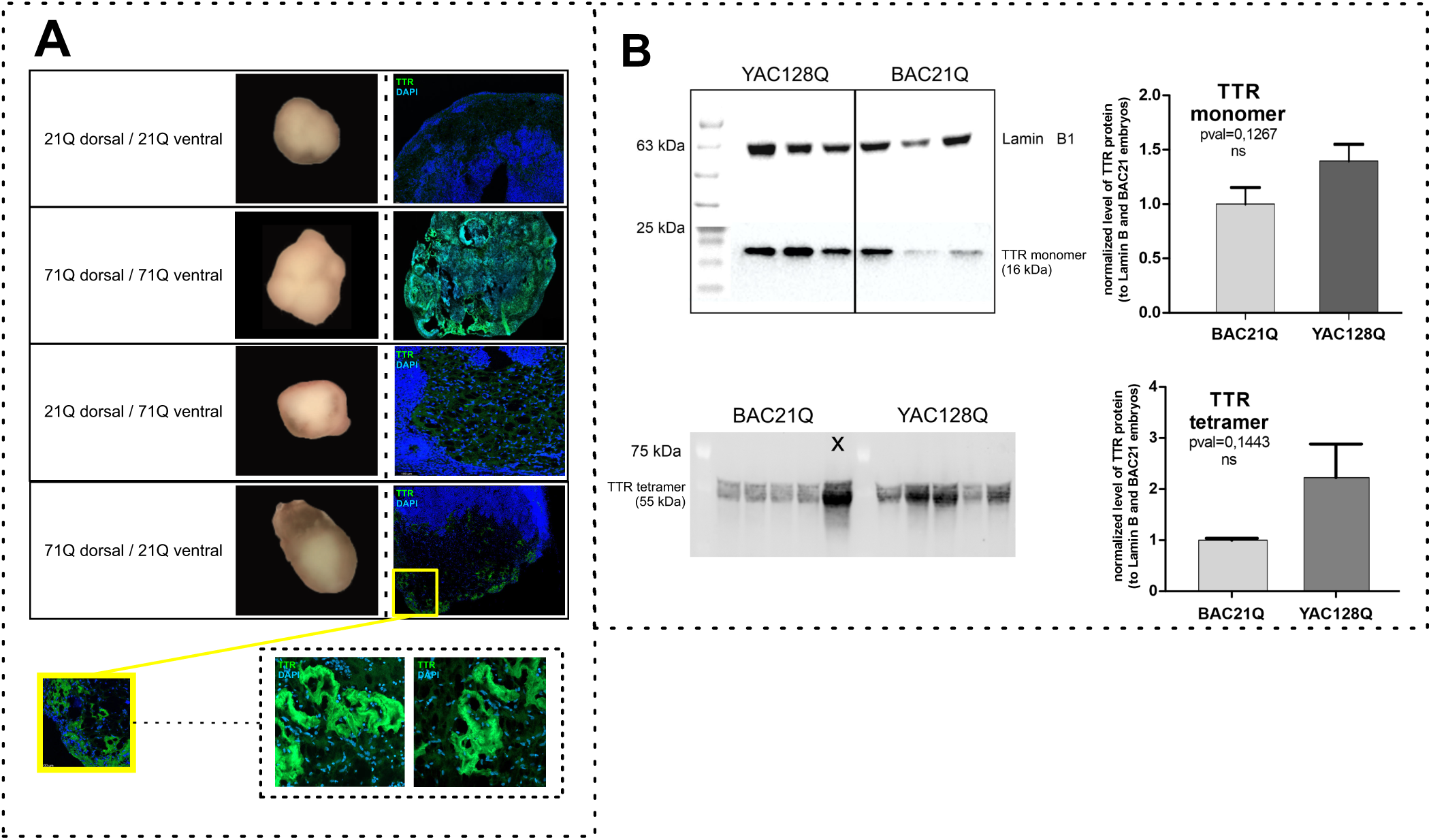
Choroid plexus and *TTR* profiles in organoids by immunofluorescence and serum by Western blotting. (A) Immunofluorescence staining for TTR in 60-day-old fused dorso-ventral 21Q/71Q mosaic organoids and 21Q dorso-ventral control organoids. IF results show an abundance of TTR in 71Q dorsal/ 21Q ventral mosaic organoids compared to the other organoids, which corresponds to the scRNA-seq results. (B) Western blot analysis of blood serum from adult 6-8-month-old mice shows levels of TTR monomer and tetramer. We showed that the markers differed between individual YAC128/null mice vs BAC21Q/null controls, mainly for TTR tetramer. When analyzed jointly, it did not reach statistical significance in YAC^128Q/null^ mice compared to BAC^21Q/null^ mice. The level of TTR monomer protein (16 kDa) was corrected to the level of the reference protein LAMIN B1. The statistical test used was the independent-samples t-test (n = 3). (B) The level of TTR tetramer protein (55 kDa) was normalized to the total amount of protein in lanes stained with Ponceau S solution. The statistical test used was the independent-samples t-test (n = 5). The graphs were generated in GraphPad Prism.

## 3. DISCUSSION

Malformations or damage to the human brain during neurodevelopment can contribute to neurodegenerative disorders such as HD. Early neurodevelopmental processes that are particularly vulnerable to perturbations in HTT function include aberrant neural stem cell proliferation, neuronal differentiation, ciliogenesis, and synapse formation (Mariani et al., 2015; McRae et al., 2017). Brain organoids provide a powerful platform for modeling these early events (Galimberti et al., 2024), especially when region-specific organoids are fused to recreate interregional interactions, such as corticostriatal connectivity (Bagley et al., 2017; Birey et al., 2017).

The developmental assembly of the corticostriatal axis relies heavily on the directed tangential migration of GABAergic interneurons originating from the MGE toward the dorsal pallium (Anderson et al., 1997). This early migration establishes the essential local circuit architecture required for functional corticostriatal projection systems. In addition, interneurons in region-specific organoids recreate cortico-striatal interactions by migrating from the ventral to the dorsal domain (Bagley et al., 2017; Birey et al., 2017). The structure that develops in close proximity to both cortex and striatum and exerts a profound influence on neurogenesis is the ChP. The ChP forms early in gestation, forming a monolayer of cells with motile cilia (Guemez-Gamboa et al., 2014; Jain et al., 2010; Pala et al., 2017), regulating the biochemical environment in which neurogenesis occurs (Bitanihirwe et al., 2022; Johansson, 2014; Johansson et al., 2013; Lun et al., 2015). The ciliary cells produce CSF and various factors, including transthyretin (TTR), retinoic acid transporters, and ionic regulators, which control progenitor proliferation, differentiation, and survival (Alshehri et al., 2020) (Moore and Iulianella, 2021). Importantly, mHTT disrupts ciliary motility and CSF flow (Keryer et al., 2011), and an enlarged lateral ventricular volume is an early HD phenotype (Hobbs et al., 2010; Jahanshahi et al., 2022). These observations suggest that early ventricular enlargement is a consequence of an important neurodevelopmental ChP-CSF pathogenic mechanism rather than a secondary neurodegenerative defect.

Using fused dorsal-ventral forebrain organoids, we identified early-rising cell populations and processes altered in HD brain development. In general, fused HD organoids grew significantly larger than control organoids. We hypothesized that the HD organoids have increased neural stem cell proliferation rather than neuronal differentiation. Indeed, the dorsal part of HD organoids mimicking the developing forebrain demonstrated a rich presence of PAX6-positive neural rosettes, indicating the abundance of neural stem cells, surrounded by TBR1-positive early-born neurons. These phenotypes align with previous reports of transcriptional immaturity in HD organoids (Conforti et al., 2018). Our compensatory mosaic organoids, in which only one half of the fused organoid carried the HD mutation, showed a marked reduction in neural stem cell rosettes, approaching the number in the control organoids. This indicates that the proliferative HD phenotype is partially reversible and influenced by non-cell-autonomous signals resulting from cooperation among cell populations in the cortex and striatum. Our findings challenge the traditional, cell-autonomous view of Huntington’s disease development.

To explore this non-cell-autonomous influence, we conducted organoid conditioning with ventral organoid medium (71V). We found that although exposure to such medium reduced the hyperproliferative growth of dorsal HD organoids (71D), it did not completely suppress their characteristic proliferation. This suggests that although soluble paracrine signals can influence HD phenotype, cell-cell contact or localized niche factors are necessary for a full rescue.

Next, to better understand the cellular events underlying these HD phenotypes, we used our fused HD brain organoids for scRNAseq. Further, we investigated the roles of the dorsal and ventral regions in HD using two models: our fused mosaic HD/Control organoid compensatory model and HD and control embryos as sources of dorsal and ventral brain regions dissected separately. Moreover, we selected the E13 embryo developmental stage, which is immediately prior to the onset of the accelerating phase of choroid plexus development in mice and is associated with cell expansion and the initiation of neurogenesis (Fame and Lehtinen, 2020).

The investigation of DEGs and ORA by pseudobulk of scRNAseq showed that juvenile HD organoids undergo an early developmental shift characterized by upregulation of genes linked to initial brain formation and stemness, and downregulation of pathways required for neuronal maturation, including mitotic spindle orientation, cell polarity, cilia function, and autophagy. Strong induction of early neurodevelopmental regulators such as *EMX2* and PAX6, and reduced expression of key ciliogenesis and spindle genes (*IFT88*, *ARL13B* **Fig. 10C**), indicating that HD organoids remain trapped in a symmetrical proliferative (Mukhtar and Taylor, 2018), immature state. Asymmetrical cell divisions are crucial for cortical layer formation, disturbed in HD organoids (di Pietro et al., 2016; Godin and Humbert, 2011; Guemez-Gamboa et al., 2014; Keryer et al., 2011; Mukhtar and Taylor, 2018; Pala et al., 2017; Park et al., 2019). HD organoids and HD embryonic mouse brains consistently show massive upregulation of *TTR*. Importantly, these developmental ORA in mosaic HD/Control organoids show restoration of maturation-related gene programs (**Supp. Fig. 1 B & C**)

We performed a high-resolution scRNAseq analysis of HD specific cell states, using JSEQ to define populations of HD organoids, mosaic organoids, and embryonic HD brains. We detected that HD organoids were highly populated by *TTR*-positive ChP cells, also characterized in our scRNAseq data by the expression of *TJP1*, the *GATA3* transcription factor, prominently active in development, and *S100B*. In contrast, we observed minimal or no ChP cells in control fused organoids. In mosaic organoids, we observed a small ChP population in the 71D/21V mosaic, whereas the population was absent in the 21D/71V mosaic. Similar to HD organoids, dorsal brain parts of YAC^128/null^ and Hu^128/21^ were also highly populated with ChP cells, while this was not the case for BAC^21/null^. Moreover, in control BAC^21/null^ embryos, we revealed a robust ventral population of MGE and STR in dorsal HD tissue. Such a signatures are predominantly attributed to MGE-derived Sox6+ interneurons migrating into the dorsal cortex and is detected by scRNAseq in the dissected dorsal part of the embryo brain (Azim et al., 2009; Batista-Brito et al., 2009; Lim et al., 2018; Shi et al., 2021) **(Supp. Fig. 4)**. In contrast, in HD embryonic models (YAC^128/null^ and Hu^128/21^), where massive ChP expansion dominates the dorsal tissue, these Sox6 positive migrating interneuron populations are virtually absent **(Supp. Fig. 4)**.

Detailed analysis revealed that expanded ChP cells in HD exhibited ectopic co-expression of both dorsal and ventral positional markers, indicating a disturbance of the epithelial cell profile characteristic of ChP. Further, within the MGE, HD tissues display downregulation of Foxp1 along with premature upregulation of the postmitotic neuron marker *Etv1*. While Foxp1 normally marks interneuron with characteristics of progenitor that have migratory potential (Pearson et al., 2020), *Etv1* acts as a gene instructing cells to slow down dividing and start to differentiate into mature neurons (Ahmed et al., 2021; Lim et al., 2018; Shi et al., 2021). Consequently, the loss of *Foxp1* coupled with premature *Etv1* activation forces these young cells to mature ahead of schedule, which can affect their migration from the MGE to the developing cortex and striatum. To summarize the findings, we constructed a conceptual model (Fig. 10D) showing that excessive ChP expansion causes both spatial occupancy of the neurogenic niche and altered paracrine signaling. Such conditions might trigger a premature Foxp1-to-Etv1 transcriptional shift in MGE progenitors, affecting interneuron migration and potentially disrupting the corticostriatal axis. Notably, MGE-derived interneurons provide the balance of excitation and inhibition in corticostriatal circuits. Therefore, their improper development could contribute to the circuit hyperexcitability and increased vulnerability seen in mature HD.

Our results from the *in vitro* and *in vivo* models indicate a clear difference between soluble paracrine signaling and physical structural support in HD pathogenesis, suggesting a two-way mechanism. Soluble secretome can modulate the proliferative tissue environment, and direct cell-cell contacts or localized extracellular matrix (ECM) niche factors are essential for preserving spatial boundaries and cell migration.

These early neurodevelopmental defects in ChP and MGE provide mechanistic insight into long-term HD clinical pathology. Ventricular enlargement in HD patients is not only a structural feature of neurodegeneration and striatal atrophy, but a consequence of pathogenic mechanism associated with HD brain development. This insight aligns with recent patient cohort data demonstrating structural ChP modifications and abnormal CSF dynamics prior to clinical onset, highlighting the ChP-CSF axis as a crucial early diagnostic biomarker and therapeutic candidate in Huntington’s disease.

Together, we introduced novel organoid models for studying HD and revealed mechanism in HD development closely linked to the HTT protein’s function. Our results establish that HD is characterized by excessive ChP, altered paracrine signaling that may be CSF-related, and delayed neuronal differentiation. Our mosaic organoids demonstrate that these early HD phenotypes can be reversible and that the pathogenic influence of mutant HTT can be modulated by the microenvironment resulting from healthy-disease interactions between cells from the cortex and the striatum, revealing a non-cell-autonomous mechanism that attenuates HD phenotypes. The implications of the findings include therapies that focus not only on lowering HTT throughout the brain but also on targeting the ChP-CSF niche, ciliary signaling, or factors secreted by healthy tissue, which offers a new candidate option for early intervention.

## 4. MATERIALS AND METHODS

All experiments were conducted in accordance with the relevant guidelines and established standards.

### 4.1. Human HD Induced-Pluripotent Stem Cells Culture

Human episomal HD and control iPSC lines were obtained from the NINDS Human Genetics Resource Center DNA and Cell Line Repository (https://catalog.coriell.org/1/ninds), from the NINDS Human Cell and Data Repository (NHCDR), and from the Cedars-Sinai Biomanufacturing Center. We used three different HD lines with 71 (ND42230), 77 (CS77iHD-77nxx) and 109 CAG (ND42224) repeats, and three control lines with 17/18 (ND38555), 21 (ND42245) and 33 CAG (ND41113) repeats. Human iPSCs were cultured in chemically defined conditions in Essential 8 medium (Life Technologies) and grown on Geltrex™ LDEV-Free, hESC-Qualified, Reduced Growth Factor Basement Membrane Matrix (Life Technologies). Cells were passaged using gentle dissociation with 0.5 mM EDTA in PBS.

### 4.2. Brain Organoid culture, differentiation and fusion

We developed the protocol for generating dorso-ventral forebrain organoids based on fusion protocol (Bagley et al., 2017; Knoblich et al., 2017) from human control and HD iPSC lines using an intermediate EZ spheres (neurospheres) culture step (Ebert et al., 2013). In brief, hiPSCs were detached with 0.5 mM EDTA in PBS, placed on a polyHEMA [Poly(2-hydroxyethyl methacrylate), Santa Cruz Biotechnology] coated dishes, and then allowed to form EZ spheres (Ebert et al., 2013) the next day. On the first day, EZ spheres were cultured in E8 medium with ROCK inhibitor (Y-2736, Tocris). On the second day, E8 medium were replaced with EZ medium (DMEM, F12, 2% B27 -vit. A, 1% penicillin/streptomycin, 0.5% Glutamax, 0.1% heparin) with growth factors, bFGF and EGF (ORF Genetics). After later mechanical passaging with McIlwain Tissue Chopper (model TC752, Campden Instruments) and obtaining the appropriate shape and diameter (>500 µm), neurospheres were incubated with small-molecule inhibitors and activators of signaling pathways for dorsal (cyclopamine A: SHH inhibitor by smoothened inhibition) and ventral forebrain patterning (WNT inhibition by IWP2 and SHH activator by SAG agonist). Subsequently, the dorsal and ventral patterned organoids were fused and differentiated on the orbital shaker. Fused dorsal-ventral HD and control brain organoids, as well as our compensatory models - mosaic fused organoids (dorsal HD/ventral control and ventral HD/dorsal control), were collected on day 60 for scRNA-seq, RT-qPCR, and immunohistochemistry analyses. Dorsal and ventral organoids from each cell line, which were cultured without the fusion step, were collected on day 60 for cortical and striatal markers assessment via qPCR. Single dorsal and ventral HD 71 and control 21 organoids were subjected to conditioning starting day 65 until day 123. Organoids were incubated with 25% conditioned media mixed with new media each day from the same genotype, in the scheme dorsal to ventral and the other way round.

### 4.3. Organoids Area Measurement

Pictures of 65-123-day-old single and fused organoids were captured with a Q-scope digital microscope model QS.80200-P (Euromex). Area measurement analysis was performed in ImageJ software using the “Freehand selection” tool. The scale was set based on the photograph of a calibration ruler added to the Q-scope kit, with a known number of pixels equaling the length of 5 mm. Graphs were created using Jamovi Software and R.

### 4.4. HD mouse tissue

The HD (Hu^128Q/21Q^, Yac^128Q/null^, Bac^21Q/null^) mouse model was maintained and bred in the Center for Advanced Technology (CAT) animal facility in Poznan. Animal experimentation and handling were approved and monitored by the Local Ethical Commission for Animal Experiments in Poznan (64/2018). For scRNA-seq analyses, pregnant female mice were sacrificed, embryos harvested, and embryonic brains dissected into cortex (dorsal part) and basal ganglia (ventral part). Blood serum for WB analyses was collected from the HD mouse model, from a typical experimental group of three or five mutant mice versus three or five control mice.

The humanized Hu^128Q/21Q^ Huntington’s disease (HD) model on the FVB/J background was developed at the Centre for Molecular Medicine and Therapeutics, part of the Faculty of Medicine at the University of British Columbia, Vancouver, Canada (Southwell et al., 2017). The Hu^128Q/21Q^ (null/null) mice were created by crossing two previously developed transgenic mouse lines, Bac^21Q^ ^(null/null)^ and YAC^128^ ^(null/null)^, which either contained a BAC chromosome with the human HTT gene encoding huntingtin with 21Q or a YAC chromosome with the same gene encoding huntingtin with 128Q. Neither line contained the mouse Htt gene (Htt null/null), but both carried human HTT genes. Crossing these two lines (Bac^21Q^ ^(null/null)^ x YAC^128^ ^(null/null)^) consistently produced embryos that included: Hu^128Q/21Q (null/null)^, YAC^128 (null/null)^, and Bac^21Q (null/null)^.

Single-cell RNAseq was performed on mouse embryos of three genotypes described above (Bac^21Q (null/null)^, YAC^128 (null/null)^, and Hu^128Q/21Q (null/null)^), having only control human HTT, only mutant human HTT, and both control and mutant human HTT respectivelly. The last two HD models differ by the presence or absence of normal HTT protein (21Q). Therefore, the model system can be used to identify changes in populations that may result from the rescuing effect of normal HTT.

### 4.5. *In Silico* Analyses of Available Brain Region Transcriptomics Data

Public scRNA-seq datasets were acquired from NCBI GEO and Broad Institute Data Portal (https://singlecell.broadinstitute.org/) to select cell population markers in different brain structures for initial expression analyses. We used mouse brain datasets including hypothalamus [HYP] (7 329 cells; GSM3562050 and GSM3562051); embryonic thalamus (E12.5) [TH] (7 365 cells; GSE122012); motor cortex [MCTX] (13 539 cells; MD704, MD705); auditory cortex [ACTX] (35 594 cells; MD717, MD720, MD21); orbitofrontal cortex [OCTX] (19 531 cells; MD716, MD717); entorhinal cortex [ECTX] (80 653 cells; MD720, MD721, MD698/699); hippocampus [HIP] (6 555 cells; MD718, MD719) and cerebellum [CER] (611 034 cells). The datasets represented both single-cell and single-nucleus approaches (∼ 800.000 cells); therefore, all mitochondrial and metabolism-related genes were excluded from the analysis based on genes from MitoCarta 2.0. Data from each mouse brain region were analyzed separately using our original JSEQ_scRNAseq pipeline designed to discover the composition of cell populations in high resolution (pipeline and manual: https://github.com/jkubis96/JSEQ_scRNAseq).

The cell populations from the high-resolution analysis were next analyzed using statistical and machine learning methods, such as a decision tree (rpart), correlation (corr), and statistical cell population comparison using the Shapiro-Wilk, T-Student, and Mann-Whitney tests with differences in the expression level of marker genes on significant level p <= 0.05. Subsequently, the specific cell population (e.g.: GABAergic neurons) was separated using quantiles 1 and 3 for high and low *HTT* gene expression in particular cells. The populations were then analyzed to obtain high expression of their specific markers and high *HTT* expression (R software with RStudio IDE). Specific qPCR primers (**Table 1**) were designed for each marker gene using MEGAX, Primer3, and NCBI primer blast tools.

**Table 1.**
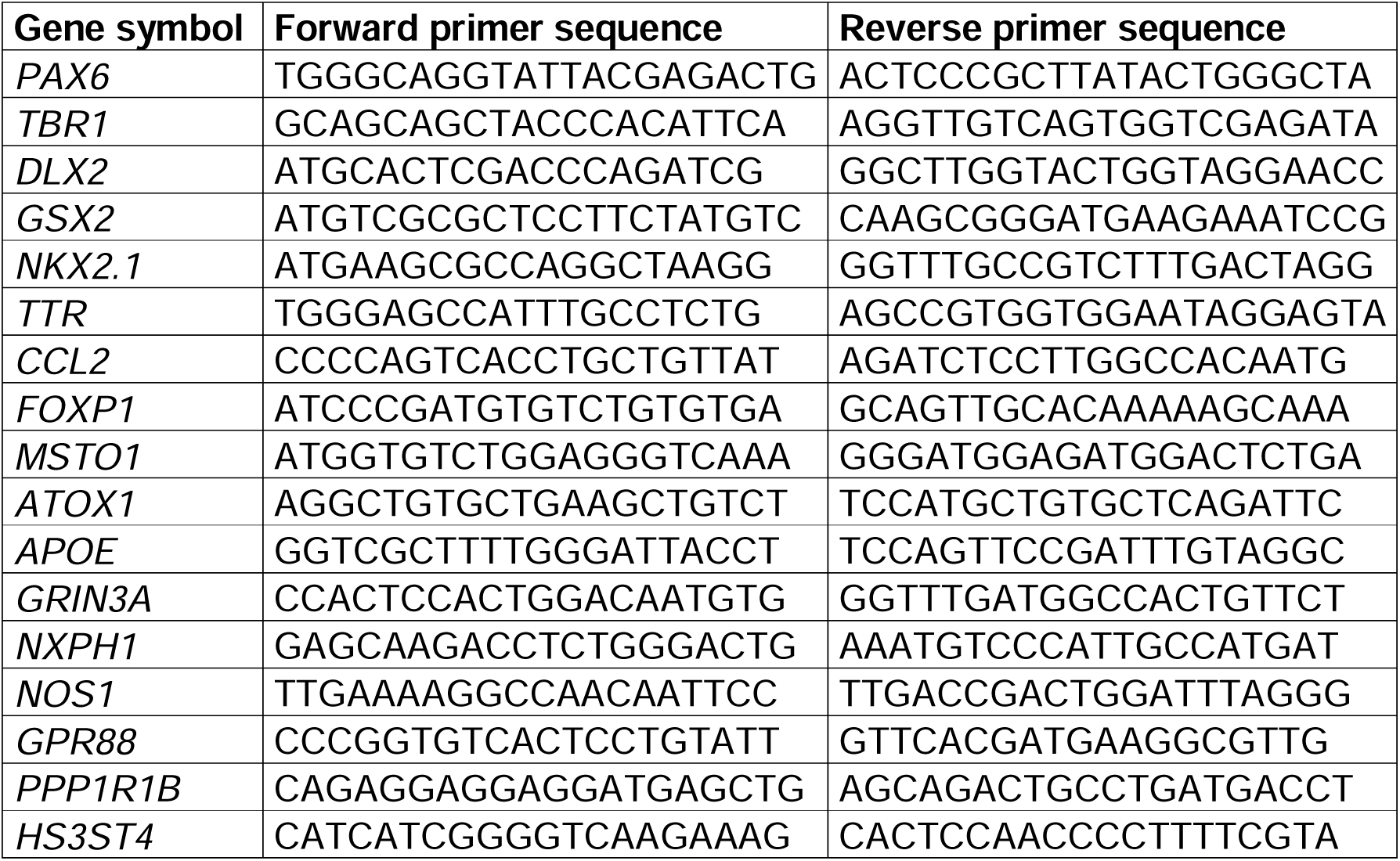
Sequences of primers used for qRT-PCR analyses.

The preanalyzed data used for this preliminary selection of markers are available and have been described (Kubiś and Figiel, 2023).

### 4.6. scRNA-seq

We performed scRNA-seq using cells from 60-day-old entire fused brain organoids or from E13.5 mouse embryonic brains dissected into dorsal (containing embryonic CTX, HIP, ChP) and ventral (containing embryonic STR, HYP, TH) parts.

In the case of organoids, our experimental setup included wild-type (WT) fused organoids 17D/17V (n = 3) and 21D/21V (n = 3); HD fused organoids 71D/71V (N = 3) and 77D/77V (N = 3); and mosaic HD and WT fused organoids 71D/21V (N = 3) and 21D/71V (N = 3).

In the case of our HD brain embryonic model, we used 3 mouse genotypes: WT Bac^21/null^, Yac^128/null^ (containing human HTT mutant allele), Hu^128/21^ (containing both mutant and WT human alleles. For each mouse genotype, we collected 1 embryonic brains (total n = 3) and further divided each brain into dorsal and ventral regions, yielding 6 independent final samples, which were subjected to scRNAseq. Summarising the scRNAseq from embryonic brains was performed from dorsal region samples Bac^21/null^ (n = 1), Hu^128/21^ ( n = 1), and Yac^128/21^ (n = 1), and the ventral region samples Bac^21/null^ (n = 1), Hu^128/21^ (n = 1), and Yac^128/21^ (n = 1).

In total, we prepared 18 libraries for organoid-derived cells and 6 libraries for embryonic mouse brains (**Fig. 1**).

The day 60 organoids and dissected dorsal and ventral embryonic brains were incubated with 0.5 ml TrypLE (Thermo Fisher Scientific) for 5 minutes at 37°C with agitation (orbital shaker; 60 RPM), followed by gentle pipette trituration, further incubation for 3 minutes, followed by another 10 times gentle trituration. TrypLE was inactivated by diluting the sample with 1 mL of PBS-EDTA containing 0.01% BSA, then centrifuging for 3 min at 300 x g. The cells were resuspended in 1 ml of PBS-EDTA with 0.01% BSA, with few pipetting actions, and passed through a 40 µm cell strainer. The cell number was determined using Trypan Blue and manual counting under a light microscope. A suspension of 300.000 cells/mL was prepared from the initial cell suspension using PBS-EDTA with 0.01% BSA.

Encapsulation of cells and further processing were prepared according to the protocol outlined in the Nadia source publication (Davey et al., 2021). In brief, the cell suspension (75,000 cells) was encapsulated with barcoded beads (150,000 beads) using the Nadia Instrument (DO BIO, UK) and disposable microfluidic cartridges, according to the pre- programmed drop-seq protocol available on the instrument. The droplet beads underwent multiple purification steps using Überstrainer and were washed with 6 x SSC, Tris-EDTA (TE), and Tween solutions to remove the oil remnants from the beads. Subsequently, the beads with captured RNA were reverse transcribed (Maxima RT Fisher Scientific, incubation 30 min RT + 90min 42°C), and the unused barcodes were removed from the beads with exonuclease I (NEB, incubation 45min 37°C), according to the Nadia protocol (Davey et al., 2021). For 3’ mRNA profiling, cDNA libraries were amplified using a PCR reaction (2,000 beads per reaction) with Kapa HiFi Mix (Roche) and Smart PCR Primer, employing the previously described protocol (Davey et al., 2021) and (Macosko et al., 2015) with modifications of the PCR cycle extending up to 15 cycles. The amplified cDNA was purified using Ampure XP beads, and concentrations were estimated using the Qubit assay. Subsequently, libraries for sequencing (NextSeq550 platform) were prepared using the Nextera XT/Vazyme kits and protocols (Illumina/Vazyme), which was modified after the tagmentation step by adding number of cycles up to 18. For library amplification, custom Index2 (I5) was used to select only tagmentation products that included UMI and BARCODE sequences (Davey et al., 2021; Macosko et al., 2015). The library size was adjusted with Ampur XP beads, and its quality and quantity was checked using a quantity control via the Qubit assay and a quality control via the TapeStation System. Sequencing was performed on the Illumina NextSeq550 platform using the NextSeq 500/550 High Output Kit v2.5 (150 Cycles). The sequencing depth was adjusted to around 10,000 reads per cell, which is sufficient for general single-cell characteristics concerning the count of organoids and mouse embryonic brains sequenced.

### 4.7. Bioinformatic Analysis of scRNA-seq data for rare cell populations

Raw FASTQ reads collected from the NextSeq 550 were analyzed using the JSEQ_scRNAseq v.2.3.2 bioinformatics pipeline. The pipeline was previously described (Kubiś and Figiel, 2023) and is available at https://github.com/jkubis96/JSEQ_scRNAseq.

In brief, the analysis in the pipeline included read quality control (FASTP) and annotation to the human GRCh38.p13 genome for organoid data, whereas for mouse embryonic brains, the data were annotated to the GRCm38.p6 mouse genome. Further pipeline steps included UMI and BARCODES quality control and repair (UMI-tools, DropSeq), data clustering with marker selection (Seurat), and other additional steps previously described in detail (Kubiś and Figiel, 2023) for the final production of high-resolution data, including rare cell populations occurring in organoids and embryonic brains. The cell populations were also annotated for their biological functions. Analyses were conducted with default settings included in the JSEQ_scRNAseq pipeline and described in the manual available on GitHub and in the publication.

Data from biological replicates were integrated within each genotype, but analyzed separately across distinct genotypes (e.g., 17D/17V, 21D/71V, 71D/71V, or embryos, e.g., HU^128/21^). This genotype-specific approach yielded higher resolution data, allowing for the precise mapping of cellular subtypes (via JSEQ_scRNAseq) to broader regional populations (e.g., CTX, STR, Ch) and enabling the identification of the ChP population. While our final analyses focus on these major cell types rather than their specific subtypes (in accordance with reviewer feedback), this high-resolution step was essential for the accurate annotation and interpretation of the complete dataset.

Downstream analyses, including data integration across genotypes (which were previously analyzed separately via JSEQ_cssg to preserve high clustering resolution - cell subtypes according to JSEQ_scRNAseq documentation), were performed using the JDtI Python library v0.2.7 (https://github.com/jkubis96/JDtI). Organoid and embryonic brain datasets were integrated independently. Cells with an unknown lineage were excluded from the analysis. During integration, the high-resolution cell subtypes identified in JSEQ_scRNAseq were merged into main cellular populations (STR, CTX, HIP, TET, ChP, MGE, CGE, LGE) to highlight broader developmental changes across these main populations.

In the JDtI workflow, Principal Component Analysis (PCA) was conducted using the top marker genes for each cell population (top 20 for organoids and top 50 for embryonic brains). These markers were selected based on a Benjamini-Hochberg adjusted p-value < 0.05 and log(FC) > 0, and were sorted primarily by Cohen’s d effect size, followed by decreasing log(FC). Using the top 25 principal components (PCs), the data were harmonized using the Harmony algorithm incorporated within JDtI. This was followed by a semi-supervised UMAP dimensionality reduction that incorporated cell population metadata. For comparison, unharmonized data and unsupervised UMAP representations are provided in the supplementary materials. UMAP parameters were set to min_dist = 0.01, n_neighbors = 15, using the Euclidean distance metric. Average gene expression was visualized using dot plots, where both dot size and color intensity reflect the average expression within a given cell population, with additional labels indicating dataset origin.

### 4.8. Identification of the differentially expressed genes (DEGs) and ORA processes

Initially, scRNA-seq transcriptomes were pooled by genotype to identify differentially expressed genes (DEGs). For organoid models, comparisons were made between HD (71D/71V and 77D/77V) and controls (17D/17V and 21D/21V), as well as between mosaic and control samples. For embryonic brains, comparisons included YAC^128Q/null^ versus BAC^21Q/null^, and Hu^128Q/21Q^ versus BAC^21Q/null^. This differential expression analysis was performed using the JDtI Python library (v0.2.7). Significant DEGs were selected based on a Mann-Whitney U test p-value ≤ 0.05 and a minimum log(FC) ≥ 0.5, and were visualized using volcano plots to compare disease versus healthy datasets.

The identified DEGs were subsequently subjected to over-representation analysis (ORA) using the GEDSpy v.2.1.3 Python library with default settings (https://github.com/jkubis96/GEDSpy). Enrichment analysis was performed for Gene Ontology (GO) terms and biological pathways (REACTOME, KEGG) utilizing a Fisher’s exact test. P-values were adjusted using the Benjamini-Hochberg false discovery rate (FDR) method, and only terms with an adjusted p-value ≤ 0.05 were considered significant. This analysis was conducted separately for up- and down-regulated gene pools.

To identify biological terms that were differentially enriched between the up- and down- regulated gene sets, we calculated a specific ’term fold change’. This metric normalizes gene occurrence and was defined as the number of term-associated genes divided by the total number of input genes in the respective pool. Using a cutoff of fold change ≥ 1.5, the qualifying terms were manually curated to retain only those associated with altered development, which were then assigned to arbitrary functional groups. These development-related core terms, initially identified from the HD and control organoid DEGs, were subsequently applied to downstream analyses of mosaic organoids and embryonic brains. The complete GEDSpy analysis output is available in the supplementary data.

### 4.9. RNA Isolation and RT-qPCR

After medium removal, HD forebrain organoids were washed once with PBS, transferred to 1,5 mL Eppendorf tubes, and then covered with 1 mL of TRIzol reagent (Thermo Fisher Scientific) according to the manufacturer’s instruction and frozen in -80°C. Upon thaw, total RNA isolation was performed according to the manufacturer’s protocol with chloroform, isopropanol, 75% ethanol, and RNase-free water. Reverse transcription was performed using Maxima H Minus Reverse Transcriptase (Thermo Fisher) (200U per reaction) on 1 µg of RNA in 20 µl of total reaction according to the manufacturer’s protocol. For priming a mixture of random hexamers (25 pmol) and oligo(dT) 18 (25 pmol) was used. Additionally, RiboLock RNase inhibitor was added to the reaction mix (20U). Before adding the enzyme and the inhibitor, templates were denatured in 65°C for 5 minutes; after mixing all reaction reagents, the reaction was incubated for 10 min at 25°C followed by 15 min at 50°C. The resulting cDNA was further diluted to a final concentration of 10 ng/uL and stored in -20°C. qPCR was performed on CFX96 Real-Time System (Bio-Rad) using HOT FIREPol® EvaGreen® qPCR Mix Plus (ROX) (SOLIS BIODYNE) on 1 µl of cDNA in 10 µl of total reaction volume. The reaction mix included 250/125 nM primers. All primers are listed in **Table 1**. Thermocycling parameters were as follows: 15 min of initial denaturation at 95°C and 39 two-step cycles with 15 s denaturation at 95°C and 60 s annealing at 60°C, followed by 5 s melt curve from 65 to 95°C, increment 0,5°C. The reaction was run on CFX96 instrument (Bio-Rad). The data were analyzed, and Cq values were determined using a regression model by CFX Manager 3.1 (Bio-Rad). ACTβ was used as a reference gene.

### 4.10. Immunohistochemistry

After medium removal, organoids were washed with PBS and fixed in cold 4% (vol/vol) paraformaldehyde (PFA) overnight at 4oC. Then, organoids were stained with Ponceau S for 20 minutes, incubated in 30% sucrose solution for 24-48 hours to protect from tissue damage during freezing, and flash-frozen in OCT CompoundTM using dry ice. Flash-frozen organoids were stored in -80°C. Organoids were sectioned using Microtome cryostat (Leica) at -20°C. 25 μm sections were collected at adhesive Superfrost PlusTM glass slides and air-dried at room temperature overnight. Then, the sections were blocked in 4% Normal Goat Serum (NGS) in TBS-T buffer for 1 h at room temperature. Primary antibodies were diluted in solution 4% NGS in TBS-T. All of primary antibodies used in this study are listed in Table 2. Secondary antibodies were used in 1:500 dilution in 4% NGS in TBS-T. Hoechst 33258 was used to visualize nuclei. A series of fluorescent confocal pictures of human brain organoids were acquired using either the TCS SP5 II (Leica Microsystems; Poland) or Opera Phoenix Plus High-Content Screening System with x40 magnification, which required assembling images along the x-, y-, and z-axis. After using the Opera Screening System, the assembly operation was conducted using JIMG v.2.1.9., available at https://github.com/jkubis96/JIMG.

**Table 2.** List of antibodies used in immunohistochemistry and/or western blot experiments.

| Antibody | Company name | Host | Diltution |
| --- | --- | --- | --- |
| TBR1 | ProteinTech | Rabbit | 1:400 |
| PAX6 | DSHB | Mouse | 1:20 |
| NOTCH1 | DSHB | Rat | 1:50 |
| NEFH | DSHB | Mouse | 1:50 |
| NES | DSHB | Mouse | 1:50 |
| MAP2 | DSHB | Mouse | 1:50 |
| GSH2 | MERCK | Rabbit | 1:500 |
| FOXG1 | Abcam | Rabbit | 1:400 |
| DLX2 | ThermoFisher Scientific | Rabbit | 1:100 |
| NKX2.1 | RD Systems | Rabbit | 1:250 |
| TTR | DSHB | Mouse | 1:50 |
| TTR | Proteintech | Rabbit | 1:1000 |
| SMI-32 | BioLegend | Mouse | 1:400 |
| LAMIN B1 | Proteintech | Mouse | 1:2000 |

### 4.11. Western Blot Analysis

Fused organoids were harvested from 60-days old cultures. Blood samples were harvested from 6-8-month-old Yac^128Q/null^ and Bac^21Q/null^ mice. All tissues were homogenized in a phosphate buffer (0.1M, pH 7.5) with 2 mM PMSF using pipetting, followed by bath sonication for 10 min. The protein concentration was estimated using a Pierce BCA protein assay kit (Thermo Scientific, Rockford, IL, USA) according to the manufacturer’s instructions. For each analysis, 20 ug of total protein was diluted in a loading buffer containing 2-mercaptoethanol and boiled for 5 min. The proteins were separated by SDS-polyacrylamide gel electrophoresis (4–15% Mini-PROTEAN® TGX™ Precast Protein Gels, BIO-RAD), transferred to nitrocellulose, and stained with Ponceau S solution. The blots were blocked with 5% non-fat milk in PBS/0.05% Tween 20 for 1 h at RT and, subsequently, incubated for 24 h at 4°C with the primary antibodies listed in **Table 2**. Lamin B1 (Proteintech) protein or bands detected after Ponceau S solution staining were used as a reference for normalization of the TTR monomer (Proteintech) or TTR tetramer (DSHB) levels in blood serum, respectively. For TTR monomer detection, the blots were probed with the respective HRP-conjugated secondary antibody (anti-rabbit or anti-mouse, 1:5000; Jackson ImmunoResearch, Suffolk, UK). The immunoreaction was detected using the ECL substrate (ThermoFisher Scientific, Waltham, MA, USA). For fluorescent Western blot analysis, a secondary antibody directed against mouse IgG, labeled with the Alexa Fluor 488 (Jackson ImmunoResearch; 1:1000) was used.

### 4.12. Statistics

Statistics were performed using GraphPad Prism 5 (version 5.00), JAMOVI 2.7.30, and R v. 4.1.2 in RStudio. In GraphPad, statistical analyses for Western blots and some qPCR analyses were conducted using t-tests to compare a single variable. In R software, analyses for qPCR and organoid size were performed. After the Levene test showed equal variance, one-way ANOVA was used for multi-group analysis, followed by post hoc t-tests (qPCR without p-value adjustment; organoid size with Bonferroni adjustment). If variance was unequal, Welch’s ANOVA was used, followed by post hoc Welch’s t-tests (qPCR without p-value adjustment; organoid size with Bonferroni adjustment). Statistical Modeling and Variance Framework.

For conditioning data, the linear mixed-effects model (LMM) was generated by JAMOVI 2.7.30 and R. Multiple organoid types were cultivated simultaneously and subjected to conditioned-medium cross-supplementation; therefore, they were analyzed as a single dataset (n = 78) to evaluate the paracrine effects and shared experimental timeframe. The LMM incorporated organoid group, conditioning, culture day, and higher-order interaction terms (group x conditioning x culture day), as fixed effects. To properly account for the longitudinal nature of the data and the correlation among repeated measurements within the same culture wells (following spontaneous aggregation into n=2), cumulative area [mm2] was measured, and individual culture wells were modeled as a random intercept. We calculated degrees of freedom and fixed-effects p-values using the Satterthwaite approximation. Group-specific estimated marginal means and 95% Wald confidence intervals were obtained directly from this combined variance. Following the multi-way ANOVA and omnibus tests of fixed effects, we performed simple slope analyses to determine the precise growth speed (in mm2 per day) for each genotype and conditioning combination across time. Simple slopes, along with their corresponding standard errors, t-statistics, and p-values, were derived directly from our LMM and the interaction.

### 4.13. Data availability

Data from single-cell sequencing of human fused brain organoids and mouse embryonic brain samples are stored on ZENOD in FASTQ (after QC) and normalized count matrix (after analysis) with metadata formats for organoids [ZENODO link] and embryonic brains [ZENODO link].

## AUTHOR CONTRIBUTIONS

K.Ś-K. generated brain organoids and performed qPCR and immunofluorescence experiments. J.K. processed the embryonic tissue and organoids for scRNAseq, performed scRNAseq library synthesis, designed scRNAseq bioinformatic tools, performed conditioning experiments, and analyzed the scRNA-seq data; M.F. and J.K. designed the scRNA-seq experiments and collected the embryonic tissue; M.F. and J.K. analyzed the conditioning experiment. M.F. performed analyses of cell positioning in organoids and embryonic brains. J.D. and B.K. generated brain organoids, M.S., Ż.K-P and P.P. collected the serum; L.H. J.P. M.R. A.S.-C. helped with scRNAseq conduction; M.H. and N.C. provided resources, the HD mouse model; K.Ś-K., J.K. and M.F. wrote the article. M.F. prepared the revision of the manuscript and performed figure editing. M.F. was responsible for the concept and obtaining funding.

## ACKNOWLEDGMENTS

This work was supported by a grant from the National Science Center (grant number 2018/31/B/NZ3/03621). We thank Professor R. Kierzek for granting access to real-time quantitative PCR machine. Cell cultures were conducted in Cell and Tissue Culture Laboratory, IBCh, PAS, Poland. Library synthesis was performed in the European Centre for Bioinformatics and Genomics (ECBIG, Poznan).

## ADDITIONAL INFORMATION

The authors declare no competing or financial interests.

## SUPPLEMENTAL DATA

**Supplementary Figure 1. ORA for upregulated and downregulated (DEGs) in brain organoids.**

The terms were arbitrarily aggregated into higher-level terms, as indicated in the red and blue boxes. The horizontal bars are named by terms (GO-, Reactome- and KEGG-terms), and the bar length represents the p-value in reverse logarithmic scale, intuitively indicating that the longer the bar, the more significant the term is. The red bars on the left indicate terms overrepresented for downregulated genes, and the blue bars on the right indicate terms overrepresented for upregulated genes. A) The ORA analysis of disease (71D/71V & 77D/77V) organoids vs. healthy (17D/17V & 21D/21V) organoids, identifying the most significant terms for upregulated and downregulated genes. This ORA analysis is the core to designate the overrepresented consensus terms, which are later used in ORA for mosaic organoids and embryonic mouse brains. B) ORA analysis of 71D/21V mosaic organoids vs. 21D/21V control organoids based on terms designated in the core analysis.

C) GSAE analysis of 21D/71V mosaic organoids vs. 21D/21V organoids based on terms designated in the core analysis. The terms that are marked as red-struck out indicate terms that are missing compared to the core analysis. In both B and C ORA, the orange bars indicate terms that are reversed in mosaic organoids compared to core (diseased vs healthy organoids) ORA analysis. The reversed and missing terms may indicate the compensation in mosaic organoids.

**Supplementary Figure 2. The ORA from upregulated and downregulated (DEGs) in dorsal and ventral embryonic mouse brain regions of YAC^128/null^, Hu^128/21^, BAC^21/null^**

The terms were arbitrarily aggregated to higher-level terms named in red and blue boxes. The horizontal bars are named by terms (GO-, Reactome- and KEGG-terms), and the bar length represents the p-value in reverse logarithmic scale, intuitively indicating that the longer the bar, the more significant the term is. The red bars on the left indicate terms overrepresented for downregulated genes, and the blue bars on the right indicate terms overrepresented for upregulated genes. The terms used in this analysis were designated in Supplementary Figure 2A (core analysis; diseased vs healthy organoids). A) The ORA analysis of diseased YAC^128/null^, vs healthy BAC^21/null^ embryonic brains. B) The ORA analysis of diseased Hu^128/21^, vs. healthy BAC^21/null^ analysis. Note that the Hu^128/21^ model contains a healthy copy of the *HTT* gene and may constitute a compensatory model compared to YAC^128/null^, which does not contain any healthy HTT copy. The orange bars in A and B indicate the terms reversed compared to the core analysis (diseased vs healthy organoids). In addition, in B, the purple bars indicate terms reversed to A (YAC^128/null^ vs BAC^21/null^).

**Supplementary Figure 3. Composition of cell population obtained from scRNAseq of HD, control, and mosaic fused dorsal/ventral organoids.**

Each panel is named by type of organoid type in the upper left corner. A) The UMAPs show cell population subtypes where each subtype is indicated by a different color. Each dot represents one cell. The UMAPs contain lists of subtypes on the right side of the panel

B) The bar plots represent the number of cells and the number of obtained subtypes in scRNA sequencing and analysis.

**Supplementary figure 4. Composition of cell population obtained in scRNAseq of YAC^128/null^, Hu^128/21^, BAC^21/null^ dorsal and ventral embryonic brain regions.**

Each panel is named by model and brain region in the upper left corner. A) The UMAPs show cell population subtypes where each subtype is indicated by a different color. Each dot represents one cell. The UMAPs contain lists of subtypes on their right side. In particular BAC^21/null^ dorsal populations representing interneurons that require intrinsic SOX6+ (Azim et al., 2009; Batista-Brito et al., 2009): MGE Sox6 Plxna4 – Cdh13; MGE Sox6 Plxna4 – Mmp16; MGE Sox6 Plxna4 – Csmd1; MGE Sox6 Plxna4 – Nnat, MGE Sox6 Plxna4 – Dcc; MGE Sox6 Plxna4 – Gm816; MGE Sox6 Plxna4 – Plxna4; MGE Sox6 Plxna4 – Dbx2; MGE Sox6 Gm10719 – Gm33838; MGE Sox6 Garem1 – Prkca; MGE Sox6 Garem1 – Gm48228; MGE Etv1 Plxna4 – Fmnl2. B) The bar plots represent the number of cells and the number of obtained subtypes in scRNA sequencing and analysis.

**Supplementary figure 5. Non-harmonized UMAP plot for scRNAseq organoid dataset.**

UMAP plots are non-harmonized equivalents of harmonized plots in the main Figure 8 A,B.

**Supplementary figure 6. Non-harmonized UMAP plot for scRNAseq organoid dataset.**

UMAP plots are non-harmonized equivalents of harmonized plots in the main Figure 9 A,B.

