## Supplementary figures and images for "Early Neurodevelopmental Defects in Huntington’s Disease are Driven by Choroid Plexus Overgrowth and Altered Paracrine Signaling"

### Supplemental Figure 1

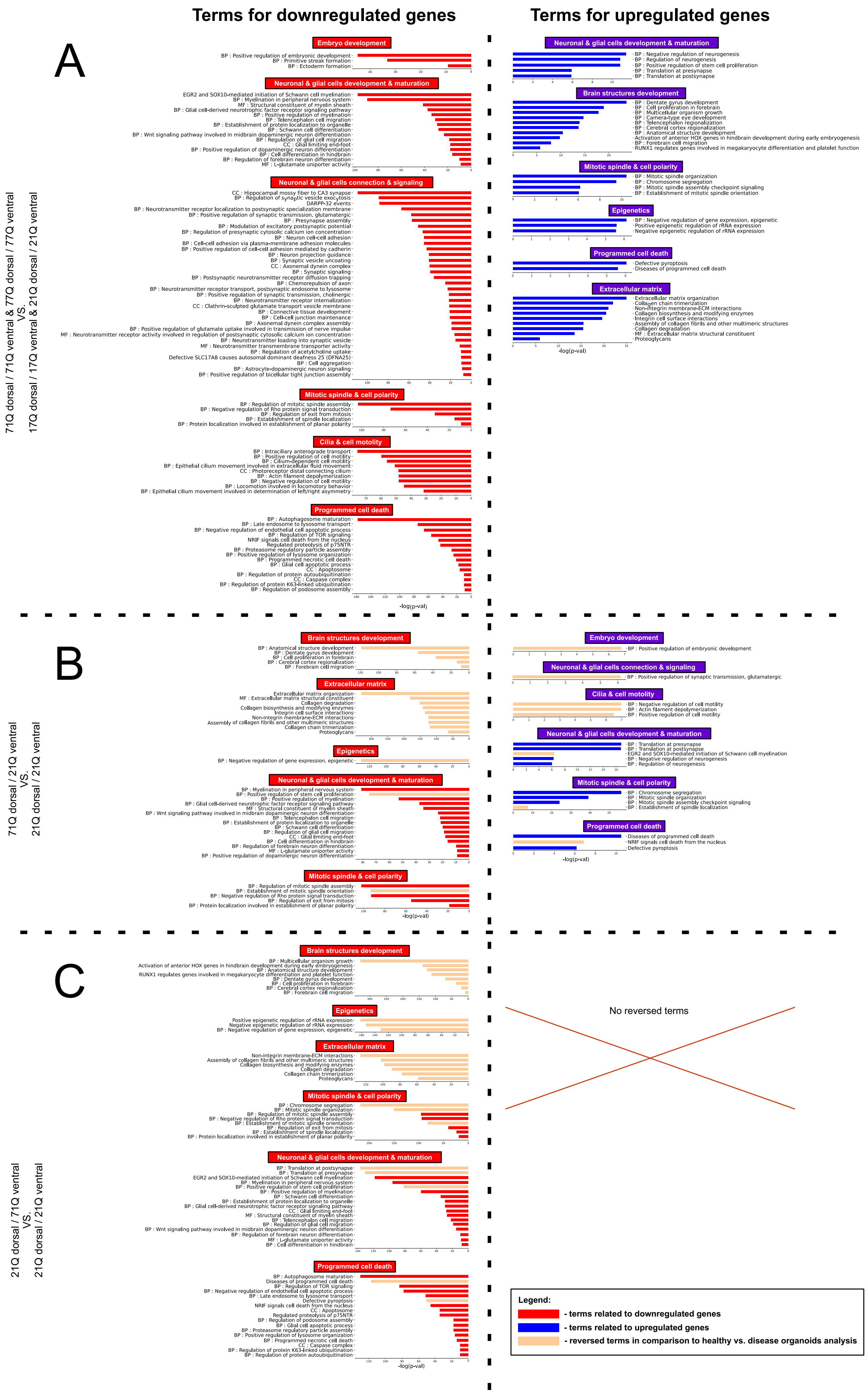

### Supplemental Figure 2

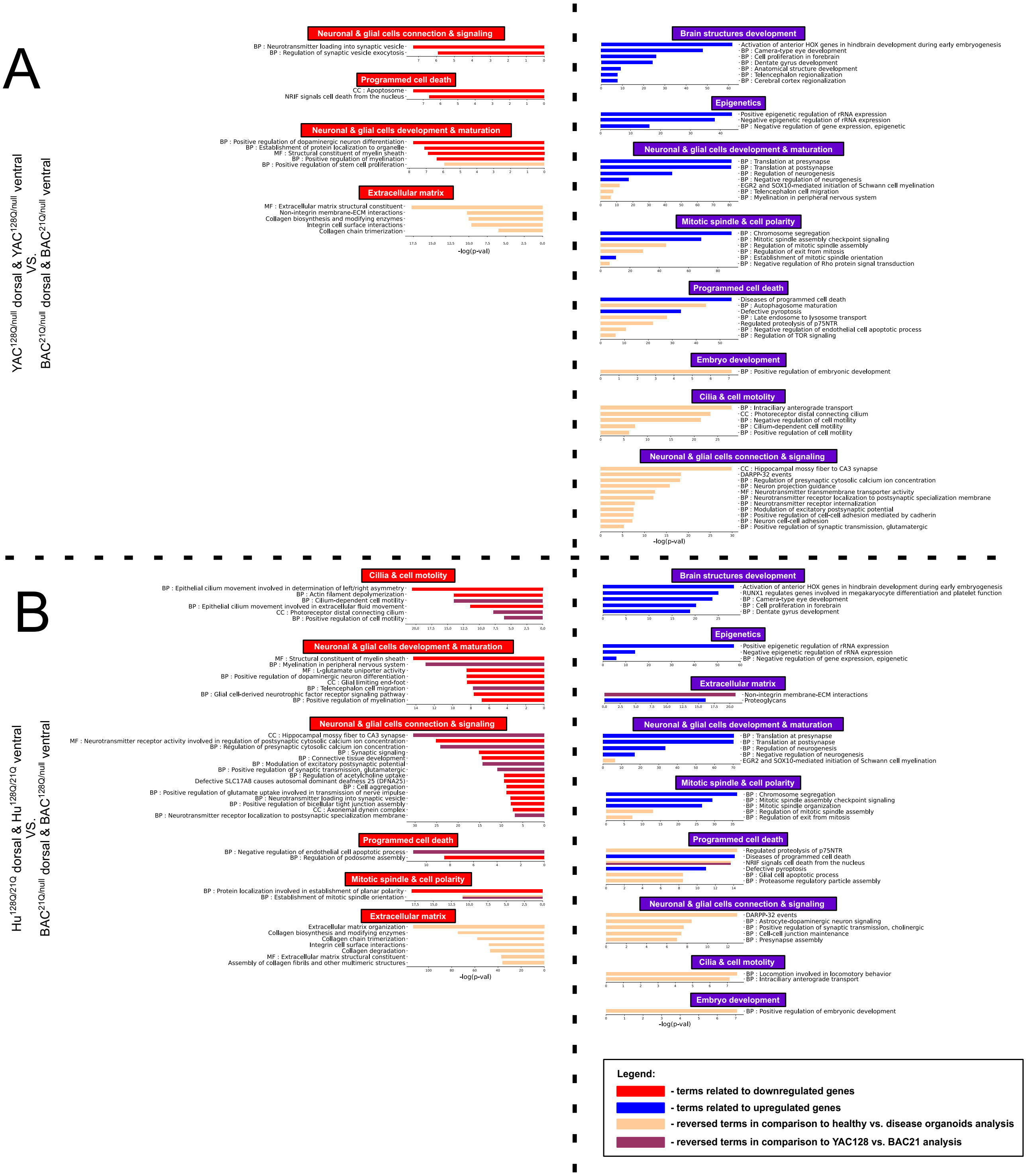

### Supplemental Figure 3

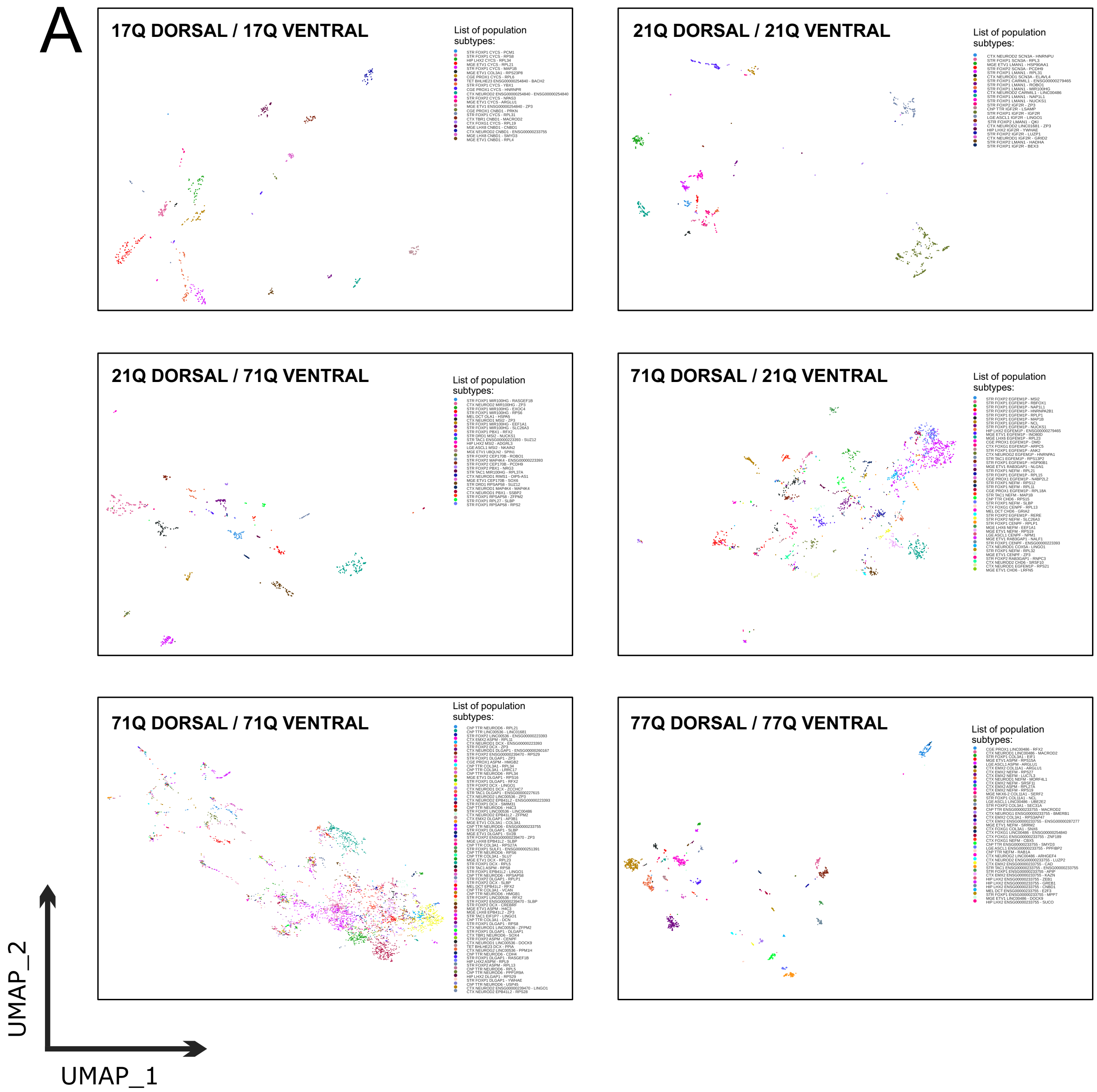

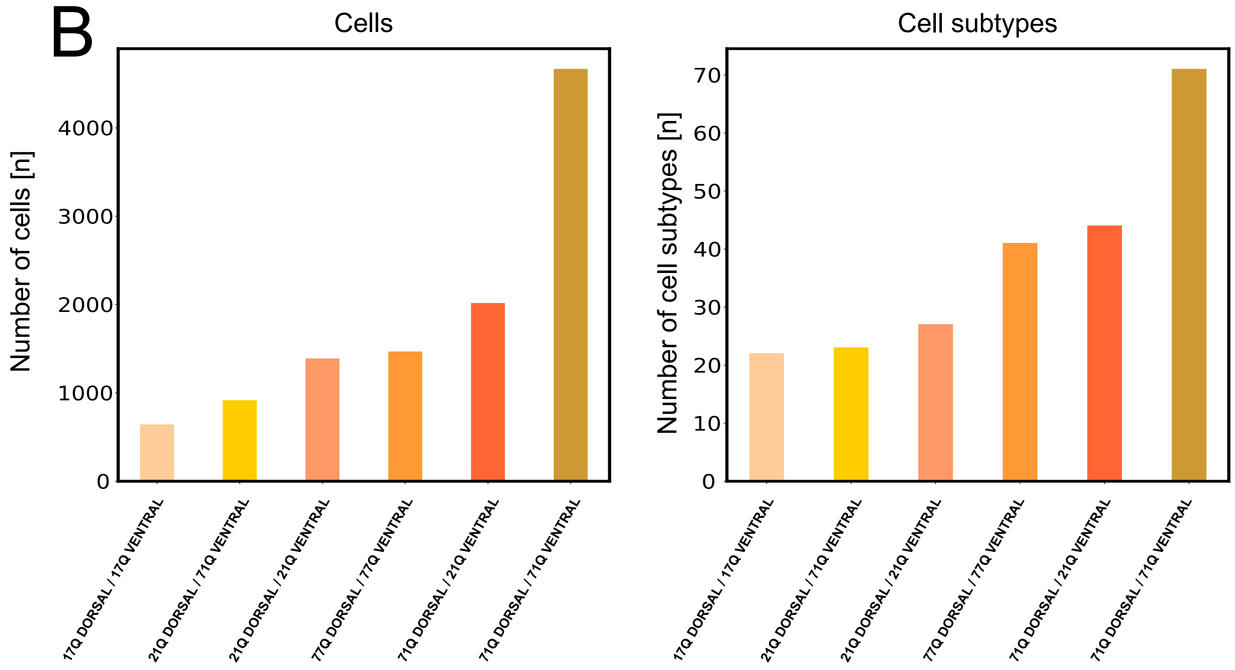

### Supplemental Figure 4

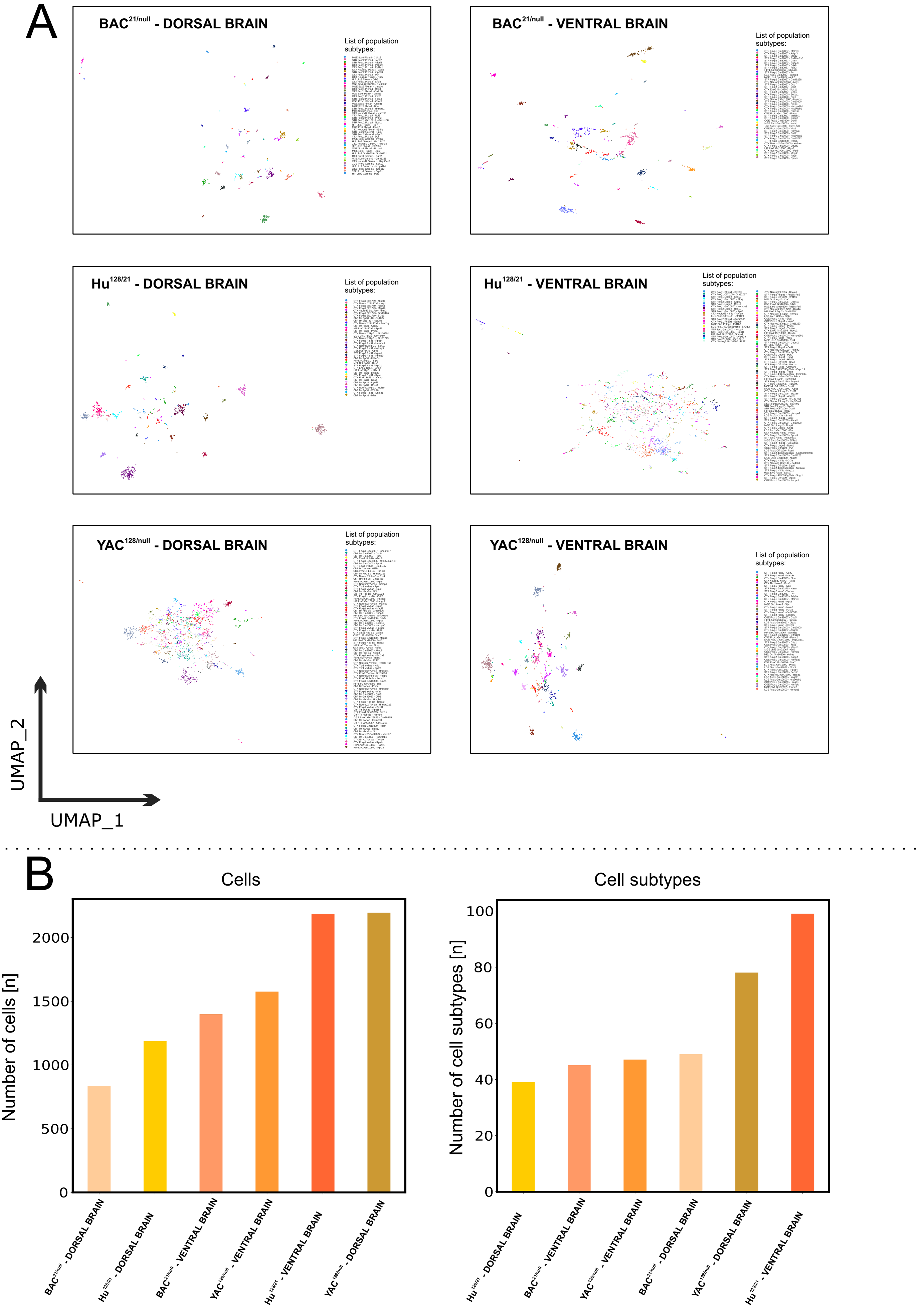

### Supplemental Figure 5

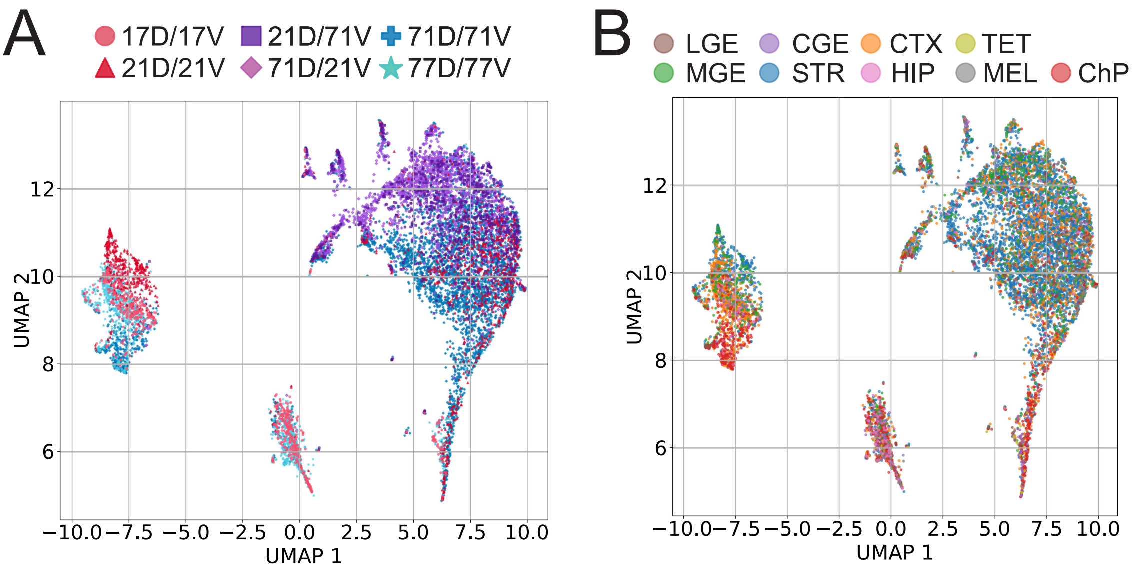

### Supplemental Figure 6

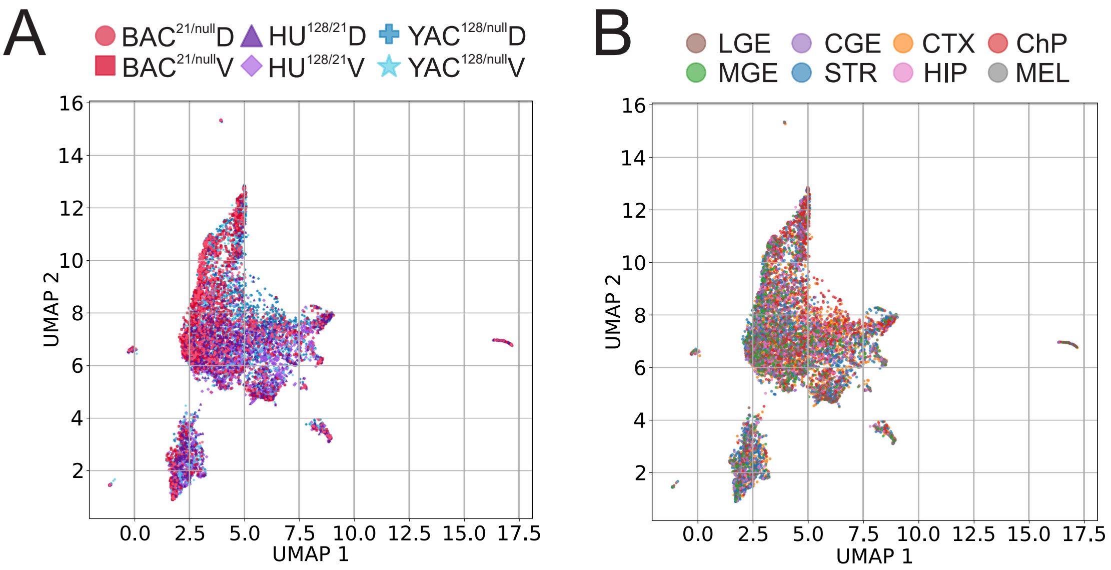
